# A combination of potently neutralizing monoclonal antibodies isolated from an Indian convalescent donor protects against the SARS-CoV-2 delta variant

**DOI:** 10.1101/2021.12.25.474152

**Authors:** Nitin Hingankar, Suprit Deshpande, Payel Das, Zaigham Abbas Rizvi, Alison Burns, Shawn Barman, Fangzhu Zhao, Mohammed Yousuf Ansari, Sohini Mukherjee, Jonathan L. Torres, Souvick Chattopadhyay, Farha Mehdi, Jyoti Sutar, Deepak Kumar Rathore, Kamal Pargai, Janmejay Singh, Sudipta Sonar, Kamini Jakhar, Sankar Bhattacharyya, Shailendra Mani, Savita Singh, Jyotsna Dandotiya, Pallavi Kshetrapal, Ramachandran Thiruvengadam, Gaurav Batra, Guruprasad Medigeshi, Andrew B. Ward, Shinjini Bhatnagar, Amit Awasthi, Devin Sok, Jayanta Bhattacharya

## Abstract

Although efficacious vaccines have significantly reduced the morbidity and mortality due to COVID-19, there remains an unmet medical need for treatment options, which monoclonal antibodies (mAbs) can potentially fill. This unmet need is exacerbated by the emergence and spread of SARS-CoV-2 variants of concern (VOCs) that have shown some resistance to vaccine responses. Here we report the isolation of two highly potently neutralizing mAbs (THSC20.HVTR04 and THSC20.HVTR26) from an Indian convalescent donor, that neutralize SARS-CoV-2 VOCs at picomolar concentrations including the delta variant (B.1.617.2). These two mAbs target non-overlapping epitopes on the receptor-binding domain (RBD) of the spike protein thereby preventing the virus attachment to its host receptor, human angiotensin converting enzyme-2 (hACE2). Furthermore, the mAb cocktail demonstrated protection against the Delta variant at low antibody doses when passively administered in the K18 hACE2 transgenic mice model, highlighting their potential as cocktail for prophylactic and therapeutic applications. Developing the capacity to rapidly discover and develop mAbs effective against highly transmissible pathogens like coronaviruses at a local level, especially in a low- and middle-income country (LMIC) such as India, will enable prompt responses to future pandemics as an important component of global pandemic preparedness.

**Highlights:**

- Identification of an Indian convalescent donor prior to emergence of SARS-CoV-2 Delta variant whose plasma demonstrated neutralization breadth across SARS-CoV-2 variants of concern (VOCs).
- Two (THSC20.HVTR04 and THSC20.HVTR26) monoclonal antibodies isolated from peripheral memory B cells potently neutralize SARS-CoV-2 VOCs: Alpha, Beta, Gamma, Delta and VOIs: Kappa and Delta Plus.
- THSC20.HVTR04 and THSC20.HVTR26 target non-competing epitopes on the receptor binding domain (RBD) and represent distinct germline lineages.
- Passive transfer of THSC20.HVTR04 and THSC20.HVTR26 mAbs demonstrated protection against Delta virus challenge in K18-hACE2 mice at low antibody doses.

**Graphical Abstract:** 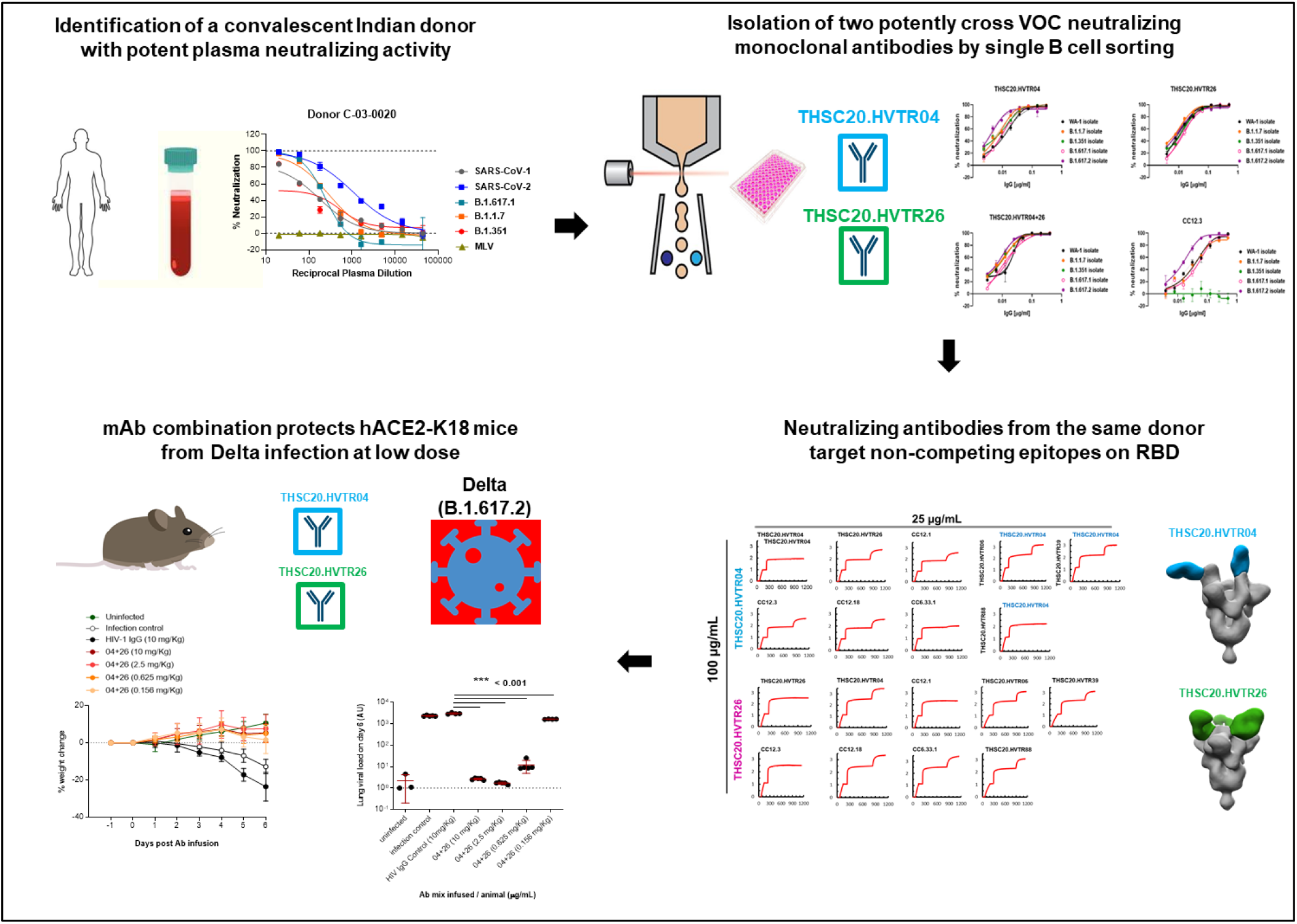

## Introduction

SARS-CoV-2 is a positive-sense single stranded RNA virus that is the etiologic cause of COVID-19, which has led to over 5 million deaths globally (https://covid19.who.int/). Although SARS-CoV-2 has a relatively low mutation rate, a combination of high transmission events (over 250 million cases to date), inequitable vaccine access, and prevailing vaccine hesitancy has led to the selection and spread of variants of concern (VOCs) that drive the persisting pandemic. These VOCs (https://www.who.int/en/activities/tracking-SARS-CoV-2-variants/) have garnered mutations that give them a selective advantage, either higher transmission, resistance to vaccine responses, or both. Among all the SARS-CoV-2 VOCs, the Delta variant (B.1.617.2), first detected in India in late 2020 in the state of Maharashtra (Dhar et al., 2021; Mlcochova et al., 2021), has become the globally dominant circulating strain and accounts for >85% sequences reported globally (https://nextstrain.org/ncov/gisaid/global). The Delta variant led to a large spike in COVID-19 cases in India as part of a second wave that culminated in over 30 million cases and over 400,000 deaths (Mlcochova et al., 2021; Thiruvengadam et al., 2021a). Since the first case of Delta was reported, scientists have determined that the Delta variant is more than twice as transmissible as the original strain of SARS-CoV-2 (Dhar et al., 2021). Some studies have also indicated that infection with the Delta variant leads to higher viral loads and worse disease prognosis compared the original Wuhan strain (Teyssou et al., 2021). Importantly, countries with high vaccination rates observed increases in cases due to Delta, indicating some breakthrough infection, but did not observe a proportional increase in hospitalization (Bergwerk et al., 2021; Lopez Bernal et al., 2021). Unfortunately, countries with limited access to vaccines face higher rates of COVID-19 cases and increases in hospitalization, which could overwhelm health systems and add to the growing tally of morbidity and mortality due to COVID-19.

Since the emergence of Delta, a new SARS-CoV-2 variant called Omicron has captured the attention of scientists and public health officials. Omicron contains a much larger number of mutations than Delta when compared to the original strain. Early studies have indicated that Omicron is four times more transmissible than the Delta variant and studies are underway on whether vaccine responses are less effective against Omicron and whether Omicron infection increases the risk for severe COVID-19 disease. As access to and uptake of efficacious vaccines remain inequitable across the globe, the likelihood of additional VOCs emerging is high, particularly among immunocompromised individuals - notably the 41% of people living with HIV (www.hiv.gov) who are not able to reliably access treatment and the millions of people around the world who are receiving treatment for cancer and other related diseases. This combination of factors leads to a perpetual unmet medical need for treatment options as global vaccine distribution and uptake languish.

In the absence of efficacious small molecule drugs at the height of the pandemic, monoclonal antibodies (mAbs) were discovered and developed as a relatively rapid countermeasure to prevent hospitalization. While effective drugs can take up to a decade or longer to discover and develop, Eli Lilly and Regeneron were able to advance mAbs from discovery to emergency use authorization (EUA) in an unprecedented 8 months. These mAbs have variable activity against different VOCs, including loss of neutralization activity of Bamlanivimab against Delta and Beta variants, loss of activity of Etesivimab against Beta and reduced activity against Alpha variant and reduced activity of Casirivimab against the Beta variant. Imdevimab, which is delivered in combination with Casirivimab as part of the REGN-CoV cocktail, remains active against Alpha, Beta, and Delta variants. The new Omicron variant, however, appears to be resistant to all four EUA antibodies (Planas et al., 2021; Wilhelm et al., 2021). To address resistant VOCs like Omicron and others that might arise, other programs have focused(Planas et al., 2021) on isolating neutralizing antibodies that target conserved epitopes on the spike protein and are therefore hypothesized to remain active against current and future VOCs. A complementary approach is to establish capacity in low- and middle-income countries (LMICs) to isolate monoclonal antibodies from convalescent donors in regions where VOCs are likely to emerge. Studies have shown that despite reduced neutralization activity against VOCs of sera from vaccinated individuals or convalescent donors infected with the original strain, those who are infected with VOCs like Beta do mount a potent neutralizing antibody response (Moyo-Gwete et al., 2021). These findings highlight the value of isolating new mAbs from convalescent donors that are effective against new VOCs to provide additional therapeutic options to prevent severe COVID-19 disease.

In the present study, we report isolation of monoclonal antibodies (mAbs) from an Indian convalescent donor by antigen-specific single B cell sorting using the receptor binding domain (RBD) of SARS-CoV-2 Wuhan strain. We report the discovery of two monoclonal antibodies (THSC20.HVTR04 and THSC20.HVTR26) that have non-competing epitope specificities on RBD and show potent neutralization of Alpha, Beta, Gamma, and Delta VOCs. Furthermore, combination of these two mAbs demonstrated significant *in vivo* protection at low antibody dose against Delta challenge in K18 hACE2 transgenic mice.

## Results

### Identification of a convalescent donor with high serum antibody titers and neutralizing activity against SARS-CoV-2 VOCs

We first screened the plasma samples obtained 6-8 weeks post infection from eleven convalescent donors in the DBT India COVID-19 Consortium cohort (Thiruvengadam et al., 2021b) who recovered from COVID-19 (Table S1). These samples were screened for anti-RBD serum titers and neutralizing activity against SARS-CoV-2 VOCs using Vero-E6 as target cells. Through this evaluation, we identified one convalescent donor (C-03-0020) whose plasma exhibited strong binding to RBD by ELISA **(Figure 1A)**, neutralization of live SARS-CoV-2 virus (Wuhan strain) (**Figure 1B**), and broad neutralization of SARS-CoV-2 (Wuhan isolate), SARS-CoV-2 VOCs, and SARS-CoV pseudoviruses **(Figure 1C)**. Based on these data, we next attempted to isolate RBD-specific monoclonal antibodies from donor C-03-0020.

**Figure 1.**
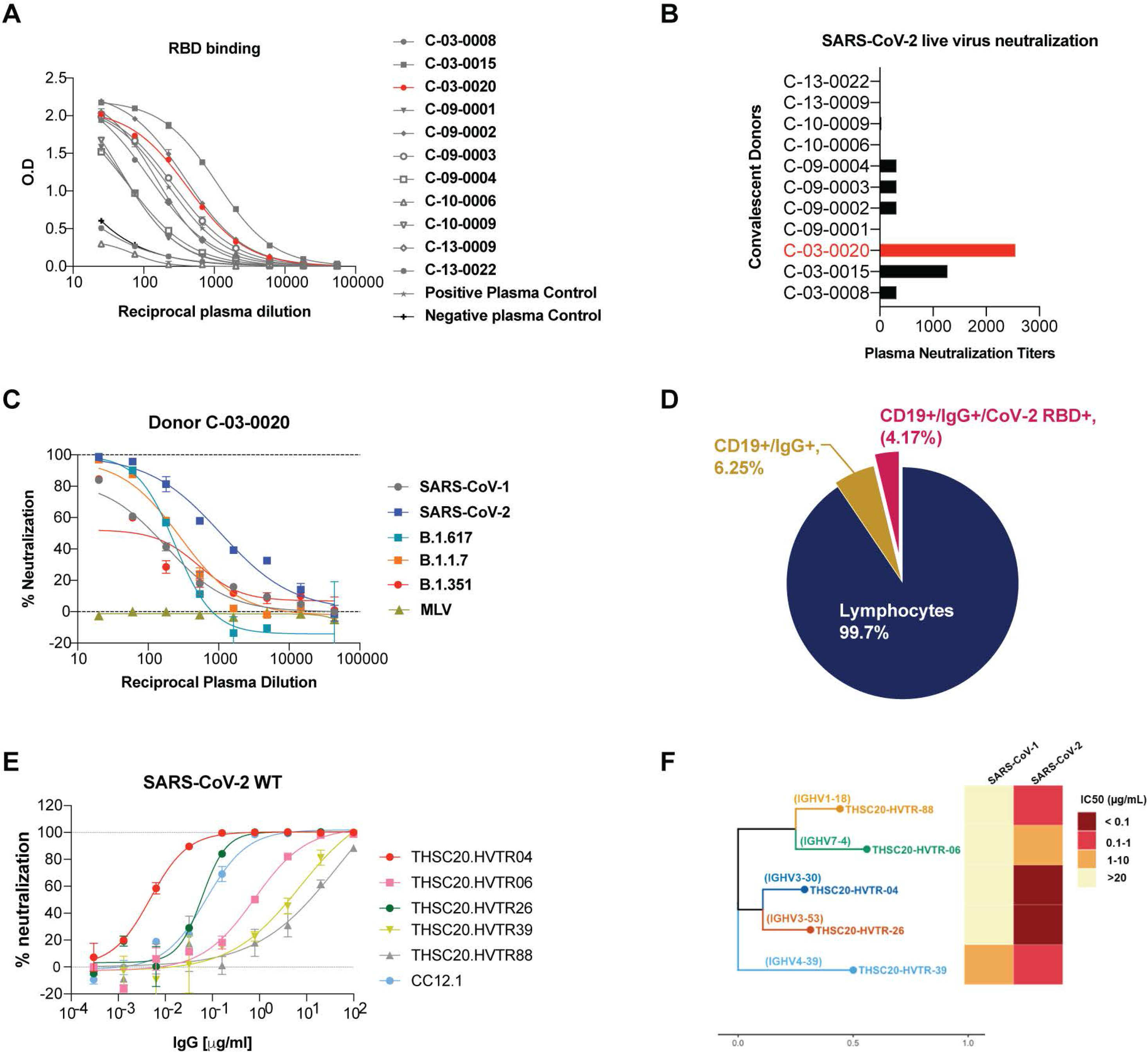
Isolation of neutralizing antibodies from the memory B cells of a convalescent donor. **A**. Identification of C-03-0020 donor with significant binding to SARS-CoV-2 RBD. **B**. High neutralizing activity of C-03-0020 plasma against SARS-CoV-2 in live virus neutralization assay. **C**. High neutralizing activity of C-03-0020 plasma against SARS-CoV-2 in pseudovirus neutralization assay. **D**. SARS-CoV-2 RBD-specific single CD19^+^/IgG^+^ B cells were isolated by FACS sorting. Percent recovery of RBD and non-RBD -specific CD19^+^ and IgG^+^_B cells are shown here. **E.** Neutralization potential of RBD-reactive mAbs were evaluated against SARS-CoV-2 pseudovirus. F. CDHR3 sequence, germline usage neutralization potency of isolated mAbs against SARS-CoV and SARS-CoV-2.

### Isolation of RBD-specific anti-SARS-CoV-2 monoclonal antibodies (mAbs)

Although anti-SARS-CoV-2 antibodies targeting different regions of the viral spike such as RBD, S1, S2 domains and N-terminal domain (NTD) have been reported (Andreano et al., 2021; Cerutti et al., 2021; Chi et al., 2020; Du et al., 2021; Graham et al., 2021; Hansen et al., 2020; Kumar et al., 2021b; Liu et al., 2020; McCallum et al., 2021; Noy-Porat et al., 2021; Rogers et al., 2020; Suryadevara et al., 2021), most of the potently neutralizing mAbs target the RBD of SARS-CoV-2, presumably because they bind to ACE2 receptor. We therefore utilized RBD as antigen bait to rapidly isolate anti-SARS-CoV-2 mAbs using single B cell FACS sorting method as described previously (Rogers et al., 2020). Using this strategy, a total of 48 SARS-CoV-2 RBD specific single B cells were antigen-sorted **(Figure S1),** mRNA from sorted cells were reverse transcribed and heavy and light chain variable regions were subsequently amplified by PCR. We successfully amplified heavy and light chains from 38 out of 48 RBD-specific B cells (79% efficiency) **(Figure 1D)**, which were then cloned into expression vectors and produced as recombinant monoclonal antibodies. Antibody transfection supernatants from these 38 heavy and light chain pairs were next screened for expression and binding to SARS-CoV-2 RBD by ELISA. As shown in **Figure S2,** 5/ 38 mAb supernatants showed expression by Fc capture ELISA and binding to SARS-CoV-2 RBD. While all the five mAbs as purified IgGs neutralized pseudovirus expressing spikes of SARS-CoV-2 wild type, Alpha, Beta and Kappa (with IC50 ranging from 0.003 - 7.2 µg/mL), one of the mAbs (THSC20.HVTR39) showed evidence of neutralizing both SARS-CoV and SARS-CoV2. (**Figure 1 & Figure S4)**.

Interestingly, sequence analysis revealed that all the five neutralizing mAbs derived from distinct germline lineages as shown in **Figure 1F and Table S2**, indicating a polyclonal neutralizing antibody response following infection in this donor. Except for THSC20.HVTR88, which has a kappa light chain, all the other four mAbs (THSC20.HVTR04, THSC20.HVTR06, THSC20.HVTR26 and THSC20.HVTR39) have lambda light chains. We also note that mAb THSC20.HVTR26 is derived from the IGHV3-53 heavy chain gene, which was reported to be a common variable heavy chain gene for SARS-CoV-2 nAbs **(Table S2)** (Yan et al., 2021). We next evaluated the level of somatic hypermutation in the variable genes. Similar to previously reported findings (Andreano et al., 2021; Pinto et al., 2020; Rogers et al., 2020; Zost et al., 2020), the isolated nAbs have relatively low levels of somatic mutations that range between 94.5% to 98.25 identical to germline. The H-CDR3 length of isolated mAbs range from 16 to 23 amino acids (aa). For the corresponding light chains, the level of IGLV gene somatic mutation ranges from 93.01 to 97.92, while the mAb with a kappa chain is 100% identical to its IGKV germline **(Table S2).** These findings counter the hypothesis that prior memory B cell responses from seasonal coronaviruses might have been responsible for the low disease severity during the first wave of infections in India, which would have resulted in recall of memory responses and higher levels of somatic hypermutation following SARS-CoV-2 infection.

### Neutralizing antibodies are active against SARS-CoV-2 VOCs

Next, we examined the ability of the newly isolated mAbs to neutralize the SARS-CoV-2 VOCs (Alpha, Beta, Gamma, and Delta) and VOIs that have circulated in India (Kappa, B.1.36 and Delta plus). First, we examined the binding affinity and avidity of the five newly isolated mAbs to the SARS-CoV-2 receptor binding domain (RBD) protein by biolayer interferometry (BLI) and ELISA respectively. As shown in **Figure 2A and Figure S3A**, BLI 8 analysis showed RBD binding affinity (KD) of these five mAbs for RBD ranging from KD values between 0.19-0.813 nM, with THSC20.HVTR04 having the highest affinity (0.19 nM) and THSC20.HVTR39 having the lowest affinity (0.813 nM). These values closely match apparent affinity measurements by ELISA to RBD with EC50 values ranging from 0.19-0.06 µg/mL **(Figure S3B)**. THSC20.HVTR04 and THSC20.HVTR26 showed the highest affinity in these assays.

**Figure 2.**
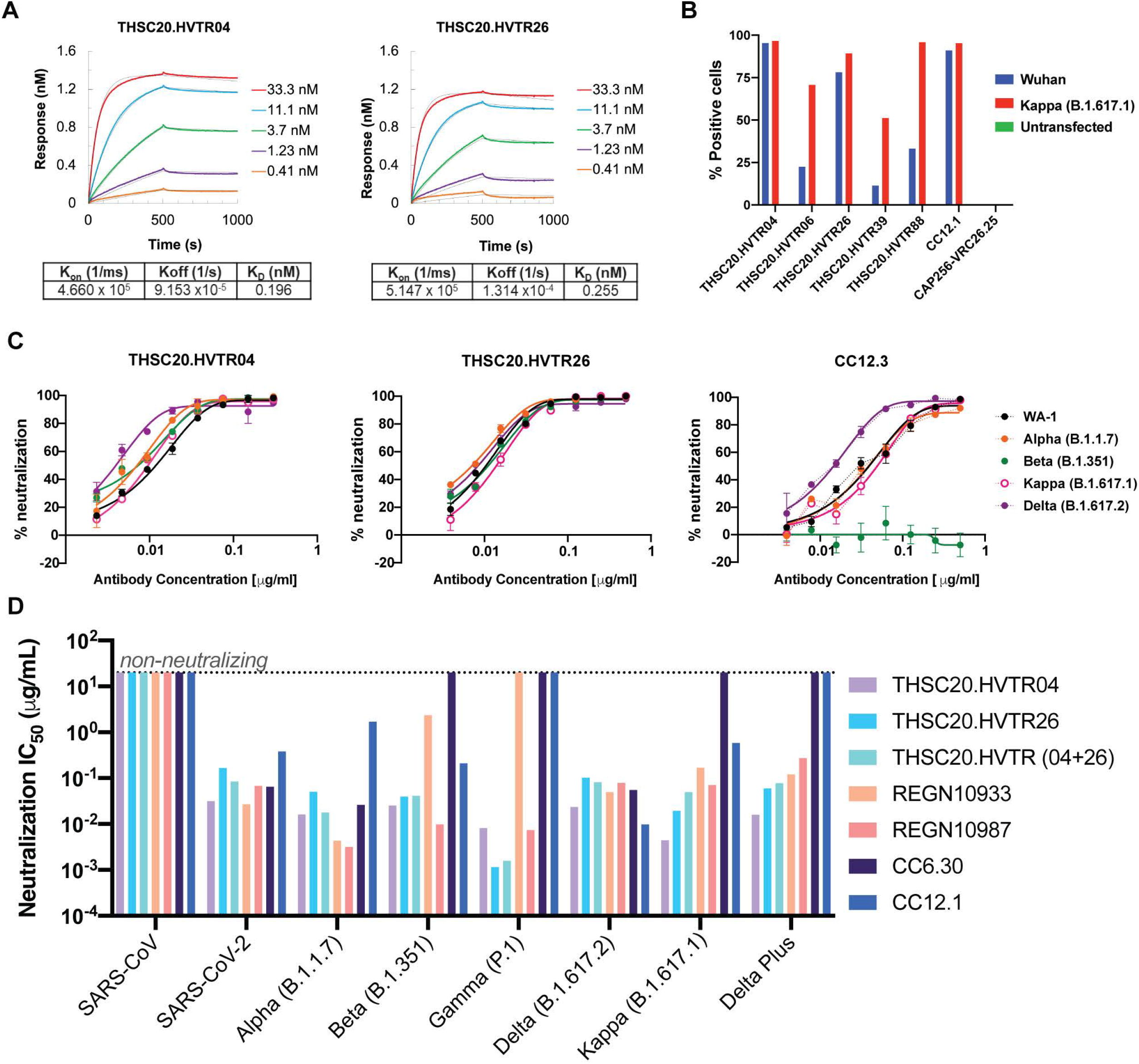
Isolated neutralizing antibodies can neutralize all circulating variants of concern. **A**. Binding affinities of THSC20.HVTR04 and THSC20.HVTR06 to the SARS-CoV-2 (Wuhan) receptor binding domain (RBD) protein by BLI-Octet. Biotinylated wild type SARS-CoV-2 RBD antigen was immobilized on Streptavidin (SA) biosensors and binding affinity of monoclonal antibodies to RBD was tested using three-fold serial dilutions of mAbs starting with 33.3 nM and lowest 0.41 nM (five different concentrations were tested). Association and dissociation was assessed for 500 seconds each. Data shown is reference- subtracted and aligned using Octet Data Analysis software v11.1 (Forte Bio). Curve fitting was performed using a 1:1 binding model and K_on_, K_off_ and K_D_ values were determined with a global fit. **B**. Binding of mAbs to SARS-CoV-2 spike protein expressed on 293T cells as assessed by mean fluorescent intensity (MFI) in a flow cytometry. **C**. Live virus focus reduction neutralization assay. The ability of the two top mAbs (THSC20.HVTR04 and THSC20.HVTR26) was assessed by dose-dependent foci reduction neutralization (FRNT) live virus neutralization assay in Vero-E6 cells. **D**. Expanded pseudovirus neutralization assay of THSC20.HVTR04 and THSC20.HVTR26 against circulating VOCs (Alpha, Beta, Gamma, Delta) and VOIs (Kappa, Delta Plus). Other known mAbs (REGN10933, REGN10987, CC12.1 and CC6.36) were included in the experiment as benchmarking controls. Representative dose response curves are shown with each concentration response tested in duplicate. Values shown are mean with SEM.

Since we used monomeric RBD to isolate the mAbs, we further assessed the ability of the isolated mAbs to bind to trimeric spike protein expressed on the cell surface. For this purpose, we expressed the SARS-CoV-2 spikes representing Wuhan wild-type or Kappa (B.1.617.1) variant on the surface of 293T cells and measured binding of mAbs to cell surface spike by fluorescence-activated cell sorting (FACS). As shown in **Figure 2B**, THSC20.HVTR04 and THSC20.HVTR26 bound more strongly than the other mAbs to the spike proteins expressed on the surface of 293T cells.

We next evaluated all of the mAbs for neutralization activity against SARS-CoV-2 variants in a pseudovirus assay. The panel of SARS-CoV-2 variants included the Wuhan strain, Alpha, Beta and Kappa strains. We found that THSC20.HVTR04 and THSC20.HVTR26 neutralized these SARS-CoV-2 variants with the highest potency (**Figure S4**). This observation is consistent with their high binding affinity to RBD as shown above. Additionally, we observed that mAb THSC20.HVTR39 neutralized SARS-CoV with IC50 of 2.9 µg/mL, indicating that this particular mAb is capable of neutralizing both SARS-CoV and SARS-CoV-2.

Since, THSC20.HVTR04 and THSC20.HVTR26 showed the greatest potency out of all the isolated mAbs, these were subsequently evaluated for neutralization individually and in combination in live virus focus reduction neutralization assay against the following VOC isolates: Alpha, Beta, Delta and VOI Kappa (as shown in **Figure 2C and Figure S5**). We also compared these antibodies to previously reported neutralizing mAbs against in a pseudovirus neutralization assay (**Figure 2D**). As shown in **Figure 2C and Table S3**, both THSC20.HVTR04 and THSC20.HVTR26 mAbs showed very potent neutralization of all the VOCs and the Kappa variant with IC50s ranging from 0.003-0.01 µg/mL when assessed against live virus isolates. Comparable neutralization potencies of THSC20.HVTR04 and THSC20.HVTR26 mAbs individually and in combination were also observed when assessed in pseudovirus neutralization assay (**Figure 2D and Table S4**). Both THSC20.HVTR04 and THSC20.HVTR26 mAbs potently neutralize Gamma (P1) and Delta-plus variants in the pseudovirus neutralization assay, which are resistant to some of the previously reported neutralizing mAbs. We note the modest superiority of these two novel mAbs over REGN10933 and REGN10987 mAbs when compared head to head in the same pseudovirus neutralization assay. Taken together, our data report the discovery of two highly potent mAbs (THSC20.HVTR04 and THSC20.HVTR26) from a single convalescent donor that not only bind to RBD with high affinities but also are capable of potently neutralizing all the VOCs and VOIs tested in this study.

### Neutralizing antibodies from the same donor target non-competing epitopes on RBD

To assess the epitope specificities of the newly isolated mAbs on SARS-CoV-2 RBD, we performed epitope binning using biolayer interferometry as described previously (Rogers et al., 2020). To achieve this, biotinylated RBD was first loaded on the streptavidin biosensor followed by binding of saturating mAb at a high concentration (100 µg/mL). Subsequently, binding of a competing mAb at lower concentration (25 µg/mL) to RBD was evaluated in the presence of saturating mAb. We also tested four previously reported anti-SARS-CoV-2 antibodies (CC12.1, CC12.3, CC12.18 and CC6.33.1) (Rogers et al., 2020) that target two distinct epitopes on RBD to inform our epitope binning. Our data revealed two distinct non-overlapping epitopes for THSC20.HVTR04 and THSC20-HVTR26 on the RBD. THSC20.HVTR06 and THSC20.HVTR39 did not compete with THSC20.HVTR-04 and THSC20-HVTR-26, indicating distinct epitope specificities for these mAbs **(Figure 3A)**. The epitope binning experiment also revealed that THSC20.HVTR-88 and THSC20.HVTR04 have similar or overlapping epitope specificities. Compared to previously reported antibodies, our experimental data show similar epitope specificities between THSC20.HVTR04 and CC6.33.1 (RBD-B) as well as THSC20.HVTR26 and CC12.1 (RBD-A).

**Figure 3.**
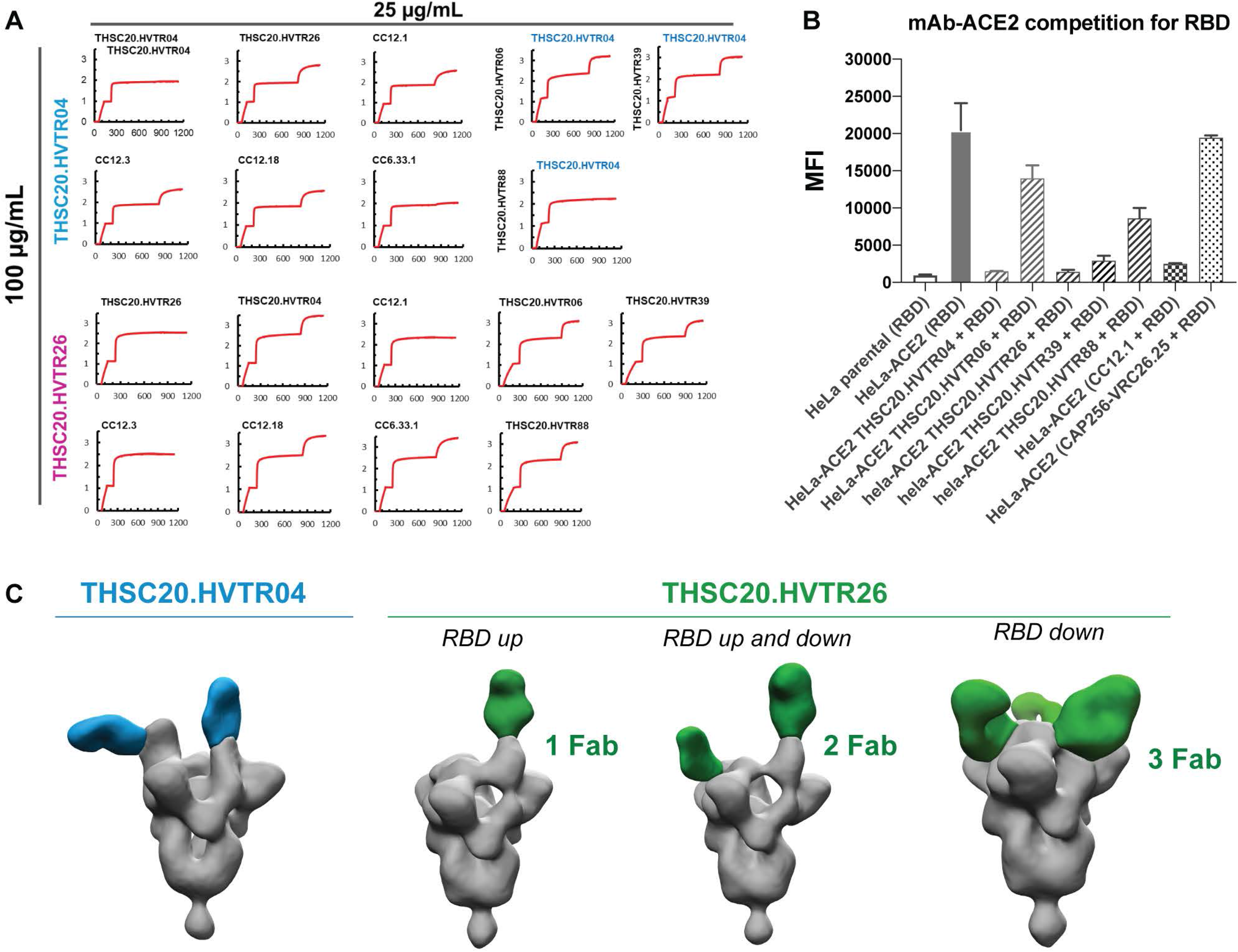
Neutralizing antibodies from the same donor target non-competing epitopes on RBD. **A.** Monoclonal antibodies were evaluated for epitope competition using BLI. Biotinylated RBD was captured using streptavidin biosensor and indicated mAbs at a concentration of 100μg/ml first incubated for 10 min followed by incubation with 25μg/ml of competing antibodies for 5 min. **B.** ACE2-mAb competition for RBD. Inhibition of SARS-CoV-2 RBD binding by five mAbs to the cell surface hACE2 was assessed by flow cytometry. **C.** Negative stain EM analysis of mAb (THSC20.HVTR04 and THSC20.HVTR26) complexed with SARS-CoV-2 spike. Data shows THSC20.HVTR04 and THSC20.HVTR26 in complex with the SARS-CoV-2 S protein.

We next examined whether the isolated monoclonal antibodies compete with RBD for ACE2 receptor binding. We carried out mAb-RBD competition for ACE2 binding assay by first incubating mAbs with RBD in a 4:1 ratio and subsequently measured binding to HeLa-ACE2 target cells by FACS. As shown in **Figure 3B**, among all the mAbs tested, THSC20.HVTR04 and THSC20.HVTR26 effectively blocked RBD binding to ACE2. We next determined which residues in the receptor binding motif (RBM) are required for antibody neutralization (Baum et al., 2020; Greaney et al., 2021; Harvey et al., 2021) for THSC20.HVTR04 and THSC20.HVTR26. We found that mutations N439K, N440K and K444N resulted in reduced sensitivity to THSC20.HVTR04, with N439K demonstrating greater than a 160-fold reduction in neutralization when assessed in pseudovirus neutralization assay (Table S4). Meanwhile, the neutralization activity of THSC20.HVTR26 was unaffected by any of these point substitutions. These data suggest that THSC20.HVTR04 might have reduced sensitivity to the Omicron variant while THSC20.HVTR26 might retain neutralization activity, both of which need to be verified experimentally.

To further characterize the epitope specificities of these antibodies, we analyzed binding of the mAbs to full length spike by negative stain electron microscopy (NS-EM). Briefly, purified IgG Fab fragments of both THSC20.HVTR04 and THSC20.HVTR26 were incubated with the trimeric spike and mAb-spike complexes were further purified by SEC as described above. Analysis of 2D class averages **(Figure S6)** and further 3D refinements of EM structural data indicated that THSC20.HVTR04 mAb binds to RBD in the up-conformation, whereas THSC20.HVTR26 can bind to RBD in different stoichiometries both in the “up” and “down” RBD conformation on the spike **(Figure 3C).** NS-EM data also showed full occupancy of the three RBD protomers on trimer spike for THSC20.HVTR26.

### THSC20.HVTR04 and THSC20.HVTR26 combination protects against Delta challenge in K18 hACE-2 mice

Based on their ability to neutralize all the SARS-CoV-2 variants with highest potency, we selected THSC20.HVTR04 and THSC20.HVTR26 to assess protection efficacy against Wuhan and Delta variant challenge in a K18-human ACE2 (hACE2) transgenic mice model (Winkler et al., 2020). Moreover, both these mAbs were negative in an ELISA-based polyreactivity assay with solubilized CHO membrane proteins **(Figure S7)**, further supporting their suitability for assessing their protective efficacy in the animal model. The experimental design of the efficacy assessment in K18-hACE2 transgenic mice model is shown in **Figure 4A and Figure S8A**. We first assessed the protective efficacy of single doses of these two individual mAbs against SARS-CoV-2 (Wuhan isolate) infection in K18-hACE-2 transgenic mice **(Figures 4B-D)**. A total of 28 mice were divided into six groups comprising five animals per group in the experimental arms and three animals in the Placebo group. Mice in groups 3, 4, 5 and 6 received intraperitoneal injection of 10 mg/kg of indicated antibodies (equivalent to 200 µg per animal) and were subsequently challenged intranasally with 10^5^ plaque forming units (PFUs) / mice of SARS-CoV-2 (Wuhan isolate) 24 hours after antibody administration. Prior to virus challenge, we obtained blood samples to determine serum IgG titers. The changes in body weight for all the experimental animals was measured daily until day 6 (end-point) **(Figure 4B)**. Additional clinical parameters were monitored to determine overall disease severity index at day 6-7 post challenge. All the mice were sacrificed on day 6 and lung tissues were harvested to measure lung virus load. As shown in **Figure 4B**, mice in the groups 2 (infection control) and 3 (non SARS-CoV-2 specific IgG control) exhibited significant weight loss of more than 10% on day 6 compared to those groups that received neutralizing antibodies. Moreover, mice that received neutralizing antibodies showed near undetectable viral RNA loads in their lung compared to the virus challenge group (group 2) the isotype control group (group 3) (P<0.001) **(Figure 4C)**. We observed a strong correlation (P<0.0001) between minimal weight loss and undetectable lung virus load in the mice that received neutralizing antibodies **(Figure 4D)**.

**Figure 4.**
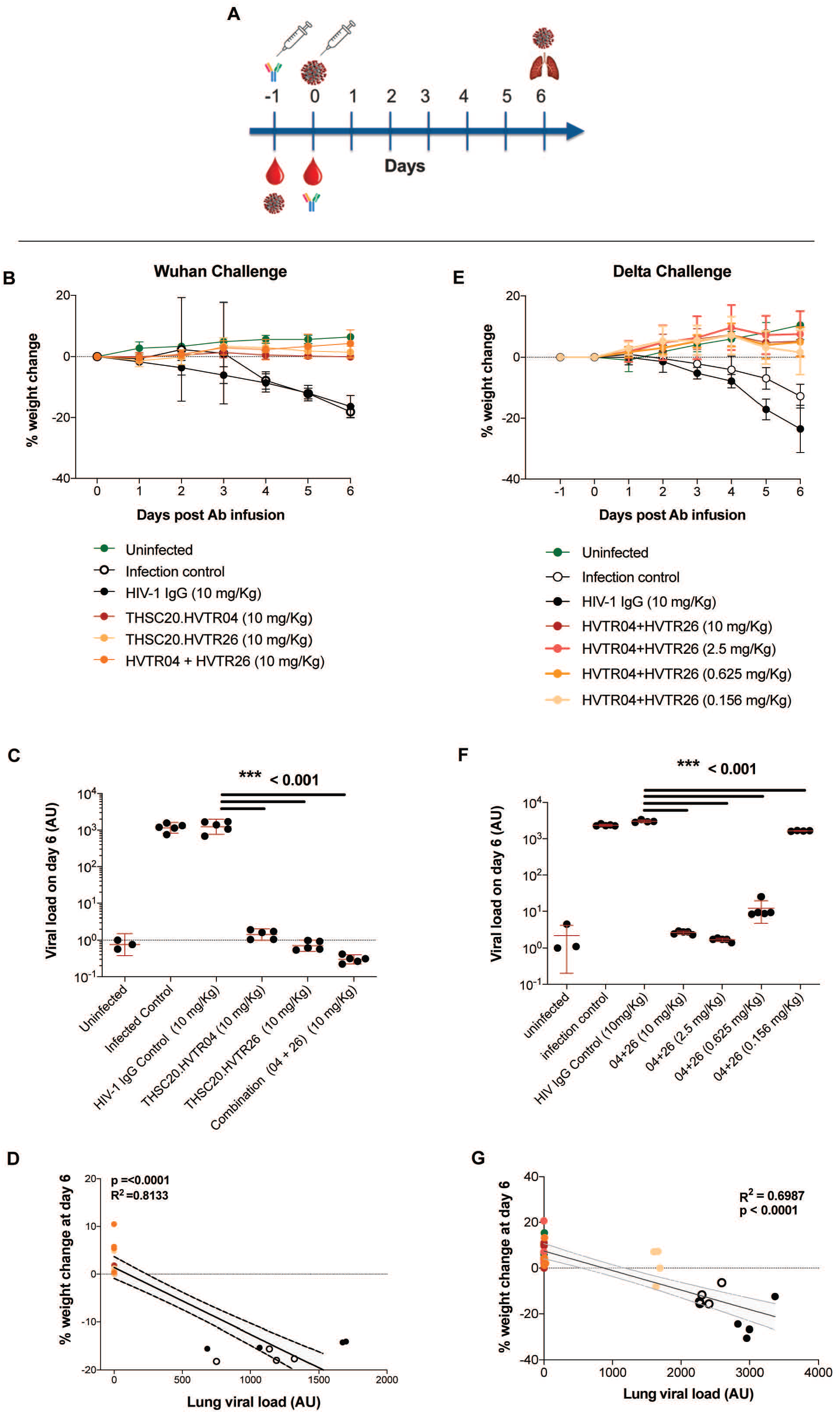
Neutralizing mAbs are able to protect against Wuhan and Delta variants in K18-hACE2 mice model. **A.** Experimental design of the efficacy assessment in K18-hACE2 transgenic mice model. K18-hACE2 mice were passively given intraperitoneal injection of the two individual mAbs or mixture of two mAbs 24 h prior to intranasal inoculation with 10^5^ PFU of SARS-CoV-2 Wuhan or Delta isolate. Each mouse was administered with single dose of 10 mg/kg body weight (equivalent to 200 ug /animal) for testing against Wuhan isolate and four different doses of 10 mg, 2.5 mg, 0.625 mg and 0.156 mg mAb per kg body weight (equivalent to 200µg, 50µg, 12.5µg and 3.125µg per animal group respectively) for testing against Delta variant isolate. **B-D.** The prophylactic effect of THSC20.HVTR04 and THSC20.HVTR26 alone and in combination against Wuhan isolate on preventing body weight loss at concentration of 10 mg per kg body weight (**B**), lung viral load assessed at day 6 (**C**) and correlation of percent body weight change with lung viral load at day 6 (**D**). **E-G.** The prophylactic effect of THSC20.HVTR04 and THSC20.HVTR26 in combination against SARS-CoV-2 Delta variant on preventing body weight loss at four different concentrations as 10 mg, 2.5 mg, 0.625 mg and 0.156 mg mAb per kg body weight, day-wise and on day 6, respectively (**E**), Lung viral load assessed at day 6 (**F**), and correlation of percent body weight change with lung viral load at day 6 (**G**).

To further assess whether these two mAbs could also demonstrate protection against the more virulent Delta (B.1.617.2) variant, we repeated the same strategy as described above. For this, we arranged the mice into seven groups **(Figure S8D)** and titrated the combination of neutralizing antibodies from 10 mg/kg (equivalent to 200 µg per animal) to 0.156 mg/kg body weight (equivalent to 3.125µg per animal) through four-fold serial dilutions. The results are comparable to the Wuhan challenge experiment **(Figures 4E-G and Figure S8)** - all the mice in the infection control group (group 2) and the isotype control group (group 3) exhibited significant weight loss at day 6, whereas no significant weight loss was observed in animals those received the antibody cocktail including those receiving the lowest dose (0.156 mg/kg body weight; group 7) **(Figure 4E)**. In addition, as an indicator of clinical prognosis, we measured lung viral RNA at day 6 and found significant reduction in the lung viral load even at an antibody dose as low as 0.625mg/kg body weight (P<0.001) **(Figure 4F)**. Interestingly, we observed near 2-fold lower lung virus load in animals those received the lowest dose of the mAb cocktail (0.156 mg/kg body weight; group 7). As expected, a strong correlation between minimal or no loss in body weight and undetectable lung virus load in the mice that received different doses of mAb combination was observed **(Figure 4G and Figure S8F)**. Taken together our study demonstrated protection by a combination of these two potently neutralizing mAbs against the highly virulent Delta variant in a transgenic hACE-2 mice model.

## Discussion

Establishing antibody discovery capacity in low and middle-income countries (LMICs) is a key component of pandemic preparedness to enable prompt responses to emerging global health challenges. The likelihood of new virus variants emerging in LMIC is high as access to vaccines and treatment options are delayed or limited and where there is an abundance of comorbidities that could accelerate the selection of new VOCs. Hence, enabling the discovery and development of monoclonal antibodies at the local level will provide new countermeasures that can be developed as treatment or prevention options that meet local needs while access to other interventions remain limited.

In our present study, we characterized mAbs that were isolated from an Indian convalescent donor in 2020 before the emergence of the deadly Delta variant. Among these mAbs, two (THSC20.HVTR04 and THSC20.HVTR26) showed neutralization of Alpha, Beta, Gamma and Delta VOCs and other VOIs including Kappa and Delta-Plus variants that originated in India. This pair of nAbs target non-competing epitopes on RBD and competed with RBD for binding to the ACE2 receptor on target cells. For THSC20.HVTR04, we identified residues N439, N440 and K444 within the RBM as critical for antibody neutralization. We did not observe point substitutions that affected the neutralization potency of THSC20.HVTR26 mAb. Of note, the ability of THSC20.HVTR04 to potently neutralize all the VOCs could be explained by the fact that none of them possess mutations at amino acid positions N439, N440 and K444 (Tao et al., 2021). Reduction in neutralization sensitivity of SARS-CoV-2 with N440K by THSC20.HVTR04 as observed in our present study indicate that recently emerged Omicron variant of SARS-CoV-2 may likely be resistant to neutralization by THSC20.HVTR04 but not by THSC20.HVTR26. Indeed, it will be important to precisely map and validate key epitope contacts for both THSC20.HVTR04 and THSC20.HVTR26 by structural studies. Our negative stain EM studies also suggest that both the mAbs binds at RBM of the RBD on viral spike, although distinctly. THSC20.HVTR04 binds RBD in the up conformation only while THSC20.HVTR26 binds to the RBD in both up and down conformations. Different classes of mAbs targeting RBD have been reported such as RBM-I, RBM-II, RBM-III and RBD core cluster I and cluster II (Finkelstein et al., 2021). Based on these observations and recent structural analyses of the variety of SARS-CoV-2 mAbs we can classify THSC20.HVTR04 in the RBM class I and THSC20.HVTR26 in the RBM class II (Finkelstein et al., 2021). Although we do not have high-resolution structural data in the present study, it is possible that our mAbs bind to their epitopes using an angle of approach and geometries that may be distinct than other potent mAbs in clinical use although they compete for the same epitope region.

To evaluate *in vivo* activity, we assessed the prophylactic efficacy of our two best mAbs (THSC20.HVTR04 and THSC20.HVTR26) individually as well as in combination using an established model of SARS-CoV-2 infection in hACE2-expressing K18 transgenic mice (Winkler et al., 2020). Mice injected with THSC20-HVTR-04 or THSC20-HVTR-26 mAbs one day before Wuhan and Delta virus challenge were completely protected from weight loss and exhibited significant decrease in lung viral load compared to mice that were given HIV-1 mAb CAP256.VRC26.25 as an isotype control. Strikingly, efficacy assessment against Delta variant with four different doses of combination of two mAbs demonstrated complete protection of mice against Delta variant at dose of as low as 0.625 mg/kg body weight. These observations confirmed that the two best neutralizing mAbs assessed here conferred complete protection against SARS-CoV-2 Delta infection in this mice model at low antibody doses. To the best of our knowledge, this is the first report of passive transfer of neutralizing antibodies and Delta virus challenge in a transgenic mouse model.

Our findings suggest that these two mAbs are valuable additions to the arsenal of existing potent mAbs against SARS-CoV-2 for development. The lessons learned from the unprecedented SARS-CoV-2 pandemic highlight the utility of timely response to such pandemics through discovery of effective mAbs against viral pathogens as one of the components of pandemic preparedness for combating coronaviruses and other deadly viruses. In the present scenario it is of paramount importance to promote and build capacity at LMICs to rapidly discover neutralizing mAbs against viral and other pathogens which will enable us to respond promptly to existing and future pandemics of coronaviruses or other highly transmissible viral pathogens.

## Authors’ contributions

JB, DS conceptualized the study; Sbhatnagar, PK, RT, SS, have contributed in cohort development and national COVID-19 biorepository under the DBT COVID-19 consortium and assisted with obtaining clinical data and biospecimen; NH along with JB and DS planned all the experiments; NH performed antigen specific single B cell sorting from donor PBMCs by FACS and amplification of antibody heavy and light chain genes with the help of SD and DR. NH, SD, PD carried out subsequent experiments that includes cloning, 14 transfection to produce mAbs, screening for expression and RBD-specific mAbs towards isolation of mAbs reported in this paper; DKR helped with Flow cytometry analysis; JS, SD, NH analyzed mAb sequences; JS performed data and statistical; NH also carried out epitope binning analysis using BLI, carried our cell surface binding assay and antibody affinity experiments using BLI; MYA, SM and NH prepared scaled up purified IgG used in all the major experiments; NH, SD, AB, SB, FZ carried out pseudovirus neutralization; SD, FZ, AB prepared different SARS-CoV-2 variant (VOC and VOI) spike constructs by side-directed mutagenesis for PSV neutralization assay; AB and FZ carried out Ab polyreactivity assay; JLT and ABW helped with the NS-EM experiment; SC, FM, GB carried out RBD-ELISA on human plasma, mice derived serum samples and purified IgG; GM carried out live virus FRNT assay; SBhattacharya, SM, KJ, SSonar contributed in live virus qualitative microneutralization screening assay; ZAR, JD and AW carried out the Ab efficacy experiments in hACE-mice model; JB, DS and NH wrote the manuscript with help from all the other authors.

## Acknowledgements.

We acknowledge the Department of Biotechnology Consortium for COVID-19 Research and all the consortium partners for making this study possible. We also thank the study participants for the clinical specimens following ethical guidelines. We thank all the staff members of our laboratories, THSTI Biorepository, THSTI Data Management team for providing us with necessary support with clinical specimens, clinical data, experiments, and research reagents. We thank Mr. Manas Ranjan Tripathy and Mr. Sandeep Goswami from Immunology Core Laboratory for the hACE2 breeding and genotyping and THSTI-SAF (Small Animal Facility) for housing and maintenance of the animals. We are very grateful to Prof Pramod Kumar Garg, Executive Director, THSTI for full support and valuable inputs and guidance. We thank Dr Sweety Samal for providing SARS-CoV-2 viruses for the animal challenge experiments and Prof Lynn Morris for providing us with CAP256-VRC26.25 HIV-1 bnAb used as isotype control in the animal challenge study. The following reagent was deposited by the Centers for Disease Control and Prevention and obtained through BEI Resources, NIAID, NIH: SARS-Related Coronavirus 2, Isolate USA-WA1/2020, NR-52281. This work was funded in part by IAVI and made possible by the support of many donors, including: the Bill & Melinda Gates Foundation, the Ministry of Foreign Affairs of Denmark, Irish Aid, the Ministry of Finance of Japan in partnership with The World Bank, the Ministry of Foreign Affairs of the Netherlands, the Norwegian Agency for Development Cooperation (NORAD), the United Kingdom Department for International Development (DFID), and the United States Agency for International Development (USAID). The full list of IAVI donors is available at http://www.iavi.org. The contents of this manuscript are the responsibility of IAVI and do not necessarily reflect the views.

## Methods

### Ethics statement for use of human samples

The participants included in this study were members of DBT COVID-19 consortium cohort, organised by interdisciplinary research institutes and hospitals in the National Capital Region of India. It was coordinated by the Translational Health Science and Technology Institute. The main clinical sites were ESIC Medical College Hospital, Faridabad, and Loknayak Hospital, New Delhi. The study protocol was approved by the Institute Ethics Committees of all participating institutions. Plasma and peripheral blood mononuclear cells (PBMCs) were prepared from blood samples obtained from eleven individuals between 6-8 weeks post recovery from SARS-CoV-2 infection who were infected in April 2020.

### Animals and ethics statement

Prior to the conduct of experiments to assess protective efficacy of the novel mAbs in small animals, approvals on the protocols involving dosing and animal challenge were obtained from the institutional animal ethics committee (approval # IAEC/THSTI/159), institutional biosafety committee (approval # 324/2021) and DBT Review Committee on Genetic Manipulation (RCGM; DBT RCGM approval #: IBKP UAC: TRARDSAB0214). 6-8 weeks old K18-hACE2 transgenic mice used to test the antibody efficacy were housed and maintained at the designated small animal facility (SAF) and subsequently transferred to the Animal biosafety level-3 (ABSL-3) institutional facility for infusion with mAbs and SARS-CoV2 challenge study. The animals were maintained under 12-hour light and dark cycle and fed with standard pellet diet and water *ad libitum*.

### FACS sorting of antigen specific memory B cells

Antigen-specific single memory B cell sorting was performed in a FACS sorter (BD FACS Melody) essentially following the methods as described earlier (Rogers et al., 2020). Briefly, cryo-preserved PBMCs were first thawed at 37^°^C in a water bath and washed with an RPMI medium containing 10% fetal bovine sera (FBS) following incubation with fluorescently-labeled antibodies (BD Biosciences) against cell surface markers for CD3 (PE-Cy7); CD8 (PE-Cy7); CD14(PE-Cy7); CD16 (PE-Cy7); CD19 (BV421); CD19 (BV421); IgD (PerCP-Cy5.5); IgG (APC-H7) in addition to labelled RBD as an antigen in FACS buffer containing PBS (pH7.4), 1% FBS, and 1.0 mM EDTA on ice. Live/Dead Fixable Aqua Blue Cell Stain (Thermo Fisher Inc.) was used to stain the cells for another 10 minutes on ice as per the manufacturer’s instructions. The avi-tagged SARS-CoV-2 RBD antigen was first labeled with biotin (Avidity, BirA500) was subsequently coupled to streptavidin-PE and streptavidin-APC (BD Biosciences) by incubating the mixture at 4^°^C for 1 hour at 4:1 molar ratio. The stained cells were subsequently washed with FACS buffer to remove unbound antibodies and probe and then filtered through 70-μm cell mesh (BD Biosciences) before processed in the FACS sorter. Single antigen RBD^+^CD3^-^CD8^-^CD14^-^CD16^-^CD19^+^CD20^+^IgD^-^IgG^+^ cells were sorted and collected into individual wells of a 96-well plate pre-filled with 20 ul of lysis buffer containing reverse transcriptase buffer (Thermo Fisher), IGEPAL (Sigma), DTT and RNAseOUT (Thermo Fisher). Plates containing sorted cells were sealed, snap-frozen on dry ice and stored at -80°C until used further.

### Amplification and cloning of variable heavy and light IgG chains

cDNA Superscript III Reverse Transcription kit (Thermo Fisher) was used to prepare from sorted cells, cDNA master mix containing dNTPs, random hexamers, IgG gene-specific primers and RT enzyme was added to generate cDNA. Heavy and light-chain variable regions of IgG were amplified in independent nested PCR reactions using specific primers. First round PCR amplification was performed using HotStar Taq DNA Polymerases (Qiagen) and second round nested PCR was performed using Phusion HF DNA polymerase (Thermo Fisher Inc.). Specific restriction enzyme cutting sites (heavy chain, 5′-AgeI/3′-SalI; kappa chain, 5′-AgeI/3′-BsiWI; and lambda chain, 5′-AgeI/3′- XhoI) were introduced in the second round PCR primers in order to clone into the respective expression vectors. Amplified PCR products were verified on the agarose gel and wells with double positives (with amplification of both Heavy and Light chain variable regions from the same well) were identified and selected for subsequent cloning experiments. PCR products were digested with specific restriction enzymes, purified and cloned in-frame into expression vectors encoding the human IgG1, Ig kappa or Ig lambda constant domains using the Quick Ligase cloning system (New England BioLabs) according to the manufacturer instructions. Ligation reactions were transformed into NEB 5-alpha competent *E. coli* cells, plated on LB agar plates containing ampicillin and incubated overnight at 37oC in the incubator. Colonies with desired inserts were screened by colony PCR and used for preparation of plasmid DNA. Plasmid clones with correct insert were further confirmed by restriction digestion with respective (New England Biolabs, Inc.) restriction enzymes before being selected for the subsequent transfection experiment. Confirmed heavy and light chain plasmid DNA were co-transfected in 293T cells (ATCC) using Fugene transfection reagent (Promega) in 24 well plates for preparing antibody supernatant for initial screening for their expression and antigen specificity as detailed in the following section. Sanger sequencing were carried out to obtain the nucleotide and amino acid sequences of variable heavy and light IgG chains. Analysis of mAb sequences were carried out using the IMGT (www.imgt.org) V-quest webserver tool.

### Capture ELISA for the detection of IgG expression

Maxisorp high protein binding 96 well ELISA plate (Nunc, Thermo Fisher Scientific.) was coated with 2μg/mL goat anti-human Fc antibody (Thermo Fisher Scientific) and incubated overnight at 4°C. Next day after washing, plates were blocked with 3% BSA in PBS (pH 7.4) for 1 hour at room temperature. After 3 times of washing with 1 X PBS containing 0.05% tween 20 (PBST), the cell supernatants which were harvested post transfection of antibody constructs in HEK 293T cells were added and incubated for 1 hour at room temperature. This was followed by addition of alkaline phosphatase-conjugated goat anti-human F(ab’)2 antibody (Thermo Fisher Scientific) Inc.) at 1:1000 dilution in 1% bovine serum albumin (BSA) incubated for an hour at room temperature. After the final wash, phosphatase substrate (Sigma-Aldrich Inc.) was added into the wells and absorption was measured at 405 nm on a 96 well microtiter plate reader.

### Streptavidin ELISA for anti SARS- CoV-2 (RBD) antibody detection

2μg/mL of Streptavidin (G-Biosciences) was coated onto each wells of Nunc maxisorp high protein-binding 96 well ELISA plate (Thermo Fisher Scientific Inc.) and incubated overnight at 4°C. Next day after washing, plates were blocked with 3% BSA in PBS (pH 7.4) for 1 hour at room temperature. 2 μg/mL of Biotinylated - RBD protein was subsequently added and incubated the plate for 2 hours at room temperature. After washing the plates for three times with PBST, cell supernatants at various dilutions were added to the wells and the plate was further incubated for 1 hour at room temperature. Finally, HRP (horseradish peroxidase) conjugated anti-human IgG Fc secondary antibody was added at a dilution of 1:1000 containing 1% BSA and the plate was incubated for an hour at room temperature. After the final wash, TMB substrate (Thermo Fisher Scientific Inc.)) was added and subsequently 1N H_2_SO_4_ was added to stop the reaction. The absorption was measured at 450 nm.

### Preparation and purification of IgG

The IgGs representing the mAbs were produced in either HEK 293T (ATCC) or Expi293 (Thermo Scientific) cells. Plasmid DNA expressing variable heavy and light IgG chains were transiently transfected into HEK293T or Expi293 cells using polyethylenimine (PEI). After 4-5 days of incubation, supernatants were harvested by centrifugation and filtered through a 0.2 μm membrane filter. Supernatants were then flowed slowly on to the Protein A (G Biosciences) beads in the column at 4°C in order to capture the secreted antibodies. Beads in the column were washed with five column volumes of 1X PBS at room temperature. Antibodies were eluted in two to three column volumes of 100 mM Glycine (pH 2.5) and immediately neutralized with 1M Tris-HCL (pH 8.0). Eluted antibodies were dialyzed using 10K MWCO SnakeSkin dialysis tubings (Thermo Fisher Scientific) against 1X PBS thrice and then concentrated in 30kDa NMWCO Amicon Ultra-15 Centrifugal Filter Units (Millipore). Antibody solutions were finally filtered through a 0.2 μm syringe filter (Thermo Fisher Scientific) before being used for further experiments. Concentration of IgG was measured by NanoDrop spectrophotometer and IgG heavy and light chain bands were visualized with 12% SDS PAGE analysis.

### Quantitative RBD-ELISA

Anti-RBD IgG ELISA was performed essentially as described in Mehdi *et al*. (Mehdi et al., 2020). For the screening of donors, plasma samples were three-fold diluted starting from 1:25 and were assessed for the presence of RBD binding IgG antibodies. To determine the mAb concentration in mice sera, a serial dilution of respective purified mAbs with known concentration was run as standard. mAb concentrations in mice sera were calculated for each sample dilution by interpolation of OD values from respective purified antibody dilutions using GraphPad Prism.

### Microneutralization screening assay

Preliminary screening of heat-inactivated plasma samples obtained from convalescent donors for their neutralization samples were serially two-fold diluted and mixed with 100 TCID_50_ of SARS-CoV-2 isolate. The virus-plasma mixture was transferred to Vero E6 monolayer seeded in 96 well plates in triplicate and incubated for 1 hour. The cell monolayer was subsequently washed with serum free media following which fresh complete medium was added. The plate was further incubated for 72 hours at 37°C in a humidified CO_2_ incubator. Absence of cytopathic effect (CPE) as an indicator of virus neutralization was assessed by observing the cells under a bright field microscope. The dilution at which no CPE was observed was considered as the neutralization titer.

### Pseudovirus (PSV) neutralization assay

Pseudoviruses expressing complete SARS-CoV2 spike genes were prepared by transient transfection of HEK293T cells with three plasmids: SARS-CoV2 MLV-gag/pol and MLV-CMV-luciferase plasmids using Fugene 6 (Promega Inc.) as described earlier (Rogers et al., 2020). After 48-hour post transfection, cell supernatants containing pseudotyped viruses were harvested and frozen at -80°C until further use. Neutralization assay was carried out using HeLa-hACE2 cells for the infection of SARS-CoV-2 wild type and variant pseudoviruses. The purified IgGs were serially diluted and incubated with pseudoviruses in a humidified Incubator at 37^0^ C. After 1-hour incubation HeLa- hACE2 cells were added to the 96-well plates at 10,000 cells/well density. After 48 hours of incubation the luciferase activity was measured by adding Britelite substrate (Perkin Elmer Inc.) according to manufacturer’s instruction and RLU obtained using a luminometer (Victor X2, Perkin Elmer Inc).

### Live virus focus-reduction neutralization test (FRNT)

The live virus neutralization assay was carried out following protocols as described by Bewley *et al*. (Bewley et al., 2021). Briefly, IgGs were serially diluted and incubated with indicated SARS-CoV-2 isolates. The virus-IgG mixtures were next added to Vero E6 cells for virus adsorption for one hour. The viral inoculum was removed, and cells were overlaid with carboxymethylcellulose and incubated for 24 hours. Cells were fixed and stained with anti- spike RBD antibody (Sino Biologicals) followed by HRP-conjugated anti-rabbit antibody (Invitrogen) and incubated with TrueBlue substrate (Sera Care). Finally, plates were washed with sterile MilliQ water, air-dried, and microplaques were quantified by AID iSPOT reader (AID GmbH, Strassberg, Germany). 50% neutralization values were calculated with four-parameter logistic regression using GraphPad Prism 7·0 software.

### Cell surface spike binding assay

The binding of mAbs to the SARS-CoV-2 spikes expressed on the HEK 293T cell-surface was assessed as described previously with some modifications (Rogers et al., 2020). Briefly, HEK293T cells were transfected with the three plasmids used to generate SARS-CoV-2 pseudovirus (SARS-CoV-2 MLV-gag/pol, MLV-CMV-luciferase and SARS-CoV-2 spike plasmids). After incubation for 36 - 48 h at 37° C, cells were trypsinized and a single cell suspension was prepared which was distributed into 96-well U bottom plates. 3-fold serial dilutions of mAbs starting at 10 µg/ml and up to 0.041 µg/mL were prepared in 50 µl/well and added to the spike expressing as well as un- transfected 293T cells for 1 hour on ice. Cells were subsequently washed twice with FACS buffer (1x PBS, 2% FBS, 1 mM EDTA) and then stained with 50 µl/well of 1:200 dilution of R-Phycoerythrin AffiniPure F(ab’)₂ Fragment Goat Anti-Human IgG, F(ab’)₂ fragment specific antibody (Jackson ImmunoResearch Inc.) for 45 min. Cells were finally stained firstly with 1 LIVE/DEAD fixable aqua dead cell stain (ThermoFisher) in the same buffer for another 15 minutes and subsequently washed twice in plates with FACS buffer. The binding of mAbs to spikes expressing on cell surface was analyzed using flow cytometry (BD Canto Analyzer). Percent (%) PE-positive cells for antigen binding were calculated and the binding data were generated. CC12.1 (SARS-CoV-2 mAb), and CAP256.VRC26.25 antibody (HIV-1 bnAb) were used as positive and negative controls respectively for this experiment.

### mAb-RBD competition assay

Inhibition of SARS-CoV-2 RBD binding by mAbs to the cell surface hACE2 was assessed by flow cytometry as described previously with some modifications (Rogers et al., 2020). Briefly, purified mAbs at 100 µg/mL and biotinylated SARS-CoV-2 RBD were mixed in 100 ul of DPBS in the molar ratio of 4:1 and incubated on ice for 1 hour. Parental HeLa and HeLa-ACE2 single cell suspension were prepared by washing cells once with DPBS and then detaching by incubation with DPBS supplemented with 5 mM EDTA. The detached HeLa and HeLa-ACE2 cell suspensions were again washed once and resuspended in FACS buffer (2% FBS and 1 mM EDTA in DPBS). 0.5 million Hela-ACE2 cells were added to the test mAb/RBD mixture and then incubated at 4°C for half an hour. 0.5 million HeLa and HeLa-ACE2 cells were incubated in separate wells with RBD alone without mAbs for use as background and positive control, respectively. After washing once with FACS buffer, HeLa and HeLa-ACE2 cells were resuspended in FACS buffer containing 1 µg/ml streptavidin-PE (BD Biosciences) and incubated for another half an hour. Cells were stained with 1:1000 final dilution of LIVE/DEAD fixable aqua dead cell stain (ThermoFisher) in the same buffer for another 15 minutes. HeLa and HeLa-ACE2 cells stained with SARS-CoV-2 RBD alone were used as background and positive control separately. The PE mean fluorescence intensity (MFI) was determined from the gate of singlet and live cells and the percentage of ACE2 binding inhibition was calculated by following formula.

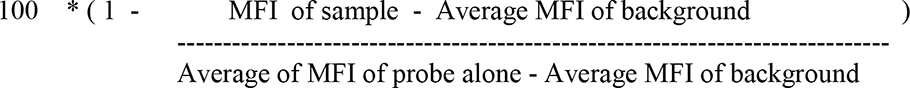

### Biolayer interferometry for assessing RBD binding affinity

Streptavidin (SA) biosensors (Forté Bio) were used to assess the binding affinities of mAbs with SARS-CoV-2 RBD in PBST (PBS containing 0.02% Tween 20) at 30 °C and 1,000 rpm. shaking on an Octet RED 98 instrument (Forté Bio Inc.). Sensors were first soaked in PBS for 15 minutes before being used to capture biotinylated SARS-CoV-2 RBD protein. RBD was loaded to the biosensors up to a level of 1.0 nm. Biosensors were then immersed into PBS for 100 seconds and then immersed into wells containing specific concentrations of a mAb dissolved in PBST (PBS containing 0.02% Tween 20) for 500 seconds to measure association. A threefold dilution series with five different concentrations (33.3, 11.1, 3.7, 1.23, and 0.41 nM) was prepared for each mAb. Biosensors were next dipped into wells containing PBST for 500 seconds to measure dissociation. Data were reference- subtracted and aligned to each other using Octet Data Analysis software v10 (Forté Bio Inc.) based on a baseline measurement. Curve fitting was performed using a 1:1 binding model and data for all the five concentrations of mAbs. Kon, Koff and *K*D values were determined with a global fit.

### Epitope binning

Epitope binning experiments were performed using Streptavidin (SA) biosensors (Forté Bio) and mAbs were binned into epitope specificities. All the incubation steps for binning experiments were performed in 1x PBS. 50-100 nM of biotinylated RBD protein antigens were loaded on streptavidin biosensors to achieve 0.9 to 1.3 nm of wavelength shift and then washed. Saturating concentration of mAbs (100μg/ml) was added for 10 min and competing mAbs at concentrations of 25 μg/ml were then added for 5 min in order to measure binding in the presence of saturating antibodies.

### Site-directed mutagenesis

Point substitutions within RBD in SARS-CoV-2 spike gene were introduced by site-directed mutagenesis using the QuikChange II kit (Agilent Technologies Inc.) following the manufacturer’s protocol and by overlapping PCR strategy as described previously (Patil et al., 2016). Successful incorporation of desired substitutions was confirmed by Sanger sequencing.

### Negative stain EM analysis

Negative stain EM (NS-EM) analysis of mAb-SARS-CoV-2 spike was essentially carried out following the protocol as described earlier (Kumar et al., 2021a). Briefly, the purified IgGs were digested for 6 hours at 37°C using 4% (w/w) liquid papain (Thermo Fischer Scientific) and digestion buffer (10 mM L-cysteine, 10X EDTA, pH 8) to prepare the Fabs. The digestion solution was then collected, and Fab fragments were purified from undigested IgG and Fc-fragments using SEC (Superdex 200 Increase; GE Healthcare). Complexes were incubated with 10–15 μg of SARS-CoV-2 spike and ∼1 mg of purified polyclonal Fab, at room temperature for 18 hours and subsequently the mAb-spike complexes were purified using SEC (Superose 6 Increase; GE Healthcare) with TBS as a running buffer and concentrated with 10 kDa cutoff Amicon ultrafiltration units. Samples were diluted in TBS and immediately deposited onto carbon-coated 400-mesh Cu grids (glow-discharged at 15 mA for 25 s), where they were then stained with 2% (w/v) uranyl formate for 30 s. Collected particle images were collected and subsequently submitted to 2D and 3D classification using Appion (Lander et al., 2009) and Relion 3.0 (Zivanov et al., 2018) data processing packages. UCSF Chimera (Pettersen et al., 2004)was used to prepare the figures by aligning representative 3D reconstructions for a specific time point and animal to each other and segmenting the maps into Fab and spike segments.

### Measuring antibody polyreactivity

The antibody polyreactivity was assessed as described earlier (Jain et al., 2017; Rogers et al., 2020). Solubilized CHO cell membrane protein (SMP) was coated onto 96-well half-area high-binding ELISA plates (Corning, 3690) at 5ug/mL in PBS overnight at 4°C. After washing, plates were blocked with PBS/3% BSA for 1 hour at room temperature (RT). Antibodies were diluted at 100ug/mL in 1% BSA with 5-fold serial dilution. Serially diluted samples were then added in plates and incubated for 1 hour at RT. After washing, alkaline phosphatase-conjugated goat anti-human IgG Fcy secondary antibody (Jackson ImmunoResearch, 109-055-008) was added in 1:1000 dilution and incubated for 1h at RT. After final wash, phosphatase substrate (Sigma-Aldrich, S0942-200TAB) was added into each well. The absorption was measured at 405 nm in a spectrophotometer. Den3 (Dengue-specific mAb) and Bococizumab (PCK9 antagonist) were included as benchmarking controls.

### K18-hACE2 mice challenge

SARS-CoV-2 Wuhan (catalogue number: USA-WA1/2020) and B.1.617.2 delta (hCoV-19/USA/PHC658/2021 catalogue number NR-55611) were procured from BEI resources (https://www.beiresources.org/) and were expanded in Vero E6 cells to produce stocks required for the experiments. Mice randomly allotted to different groups (n=5) *viz,* infection control and those received SARS-CoV-2 specific (THSC20.HVTR04 and THSC20.HVTR26) and non-specific (HIV CAP256.VRC26.25) monoclonal antibodies as IgG were housed in different cages. Antibody recipient groups were given intraperitoneal injection of IgG one day prior to challenge (day -1). Except for the unchallenged control group (n=3), animals in all other groups were challenged with 10^5^ PFU of SARS-CoV2 (Wuhan and Delta isolates) intranasally on Day 0, administered through a catheter 25 µl/ nare under anesthesia by using ketamine (150mg/kg) and xylazine (10mg/kg) inside ABSL3 facility (Chan et al., 2020; Rizvi et al., 2021b; Sia et al., 2020; Winkler et al., 2020). Unchallenged control group received mock PBS (pH 7.4) intranasally.

### Gross clinical parameters of SARS-CoV-2 infection

All the infected animals were euthanized on day 6 days post infection at the ABSL3. Changes in body weight, activity of the animals were observed on each day post challenge. Post sacrifice, lungs of the animals were excised and imaged for gross morphological changes. Right lower lobe of the lung was fixed in 10% neutral formalin solution and used for histological analysis. Rest of the lung was homogenized in 2ml Trizol solution for viral load estimation. The tissue samples in trizol were stored immediately at -80 °C till further use. Blood of the animals were drawn through retro-orbital vein on day -1 and 0 and through direct heart puncture at the end-point. Serum samples were stored at -80 °C till further use.

### Lung viral load quantification

Homogenized lung tissues were used for RNA isolation using Trizol-chloroform method as per the manufacturer’s protocol and quantitated in a Nanodrop. 1 µg of total RNA was then reverse-transcribed to cDNA using the iScript cDNA synthesis kit (Biorad; #1708891) (Roche). Diluted cDNAs (1:5) were used for qPCR by using KAPA SYBR® FAST qPCR Master Mix (5X) Universal Kit (KK4600) on Fast 7500 Dx real-time PCR system (Applied Biosystems) and the results were analyzed with SDS2.1 software. The CDC-approved SARS-CoV-2 N gene primers: 5′-GACCCCAAAATCAGCGAAAT-3′ (Forward), 5′-TCTGGTTACTGCCAGTTGAATCTG-3′ (Reverse) were used for virus load estimation. The relative expression of each gene was expressed as fold change and was calculated by subtracting the cycling threshold (Ct) value of β-actin (endogenous control gene) from the Ct value of target gene (ΔCT). Fold change was then calculated according to the previously described formula POWER (2,-ΔCT) (Rizvi et al., 2021a). For absolute quantitation, known copy number of the virus RNA was used as a standard to generate the standard curve.

## Supplementary Tables and Figures

### Supporting Tables

**Table S1.** Convalescent donor history & background information.

| **PID** | **Age** | **Gender** | **Month &**<br>**Year of**<br>**infection** | **Clinical status at**<br>**the time of blood**<br>**draw** | **Blood drawn (Days post**<br>**onset of detection)** | **Disease severity** |
| --- | --- | --- | --- | --- | --- | --- |
| C-03-0008 | 56 | Female | April, 2020 | Symptomatic | 55 | Mild/Moderate |
| C-03-0015 | 46 | Male | April, 2020 | Asymptomatic | 47 | None |
| C-03-0020 | 33 | Male | April, 2020 | Symptomatic | 57 | Mild/Moderate |
| C-09-0001 | 30 | Male | April, 2020 | Asymptomatic | 42 | None |
| C-09-0002 | 44 | Male | April, 2020 | Symptomatic | 50 | Mild/Moderate |
| C-09-0003 | 51 | Male | April, 2020 | Symptomatic | 56 | Mild/Moderate |
| C-09-0004 | 50 | Female | April, 2020 | Symptomatic | 53 | Mild/Moderate |
| C-10-0006 | 24 | Male | April, 2020 | Asymptomatic | 40 | None |
| C-10-0009 | 42 | Male | April, 2020 | Asymptomatic | 39 | None |
| C-13-0009 | 67 | Male | April, 2020 | Symptomatic | 50 | Mild/Moderate |
| C-13-0022 | 45 | Male | April, 2020 | Symptomatic | 47 | Mild/Moderate |

**Table 2.** Heavy and light chain variable IgG sequence characteristics and germline usages.

Heavy chain IgG
| mAb ID | V gene & allele | % identity to germline (V region) (nucleotide) | J gene & allele | % identity to germline (J region) (nucleotide) | CDHR3 sequence | CDHR3 length |
| --- | --- | --- | --- | --- | --- | --- |
| THSC20.HVTR04 | IGHV3-30-3*01 F | 96.88% | IGHJ2*01 F | 90.57% | CASVADTAMVPEWYFDLW | 16 |
| THSC20.HVTR06 | IGHV7-4-1*02 F | 96.53% | IGHJ6*02 F | 96.77% | CARVLGDQQVVGDDYYYYYAMDVW | 23 |
| THSC20.HVTR26 | IGHV3-53*04 F | 98.25% | IGHJ3*02 F | 96% | CARLVGALTNIVVSGDGGAFDIW | 21 |
| THSC20.HVTR39 | IGHV4-39*07 F | 94.50% | IGHJ3*02 F | 98% | CARNLFYYDRSGSSHHDAFDIW | 20 |
| THSC20.HVTR88 | IGHV1-18*01 F | 98.61% | IGHJ4*02 F | 95.83% | CAREGKYSSGWSYSEYFDYW | 18 |

Light chain IgG
| mAb ID | V gene & allele | % identity to germline (V region) (nucleotide) | J gene & allele | % identity to germline (J region) (nucleotide) | CDR3 sequence | CDHR3 length |
| --- | --- | --- | --- | --- | --- | --- |
| THSC20.HVTR04 | IGLV2-11*01 F | 97.92% | IGLJ3*02 F | 97.37% | CCSYAGSWVF | 8 |
| THSC20.HVTR06 | IGLV3-1*01 F | 97.49% | IGLJ3*02 F | 97.74% | CQAWDNSGWVF | 8 |
| THSC20.HVTR26 | IGLV2-23*02 F | 97.57% | IGLJ3*02 F | 97.22% | CCSYAGSSTWVF | 10 |
| THSC20.HVTR39 | IGLV3-21*02 F | 93.01% | IGLJ3*02 F | 100% | CQVWDRGSDVWVF | 11 |
| THSC20.HVTR88 | IGKV3-20*01 F | 100% | IGKJ1*01 F | 92.11% | CQQYGSSLWTF | 9 |

*The analysis of variable heavy (HC) and light (LC) chains IgG nucleotide sequences of the mAbs were done using by IMGT/V-QUEST tool (www.imgt.org).*

**Table S3.** Neutralization potency of THSC20.HVTR04 and THSC20.HVTR26 and their combination against replication competent SARS-CoV-2 live VOCs.

|  | IC50 (µg/mL) |  |  |  |
| --- | --- | --- | --- | --- |
|  | THSC20.HVTR04 | THSC20.HVTR26 | THSC20.HVTR 04+ 26 | CC12.3 |
| SARS-CoV-2 | 0.01 | 0.009 | 0.011 | 0.034 |
| B.1.1.7 (alpha) | 0.005 | 0.006 | 0.007 | 0.037 |
| B.1.351 (beta) | 0.006 | 0.014 | 0.007 | >20 |
| B.1.617.1 (kappa) | 0.008 | 0.011 | 0.011 | 0.044 |
| B.1.617.2 (delta) | 0.003 | 0.007 | 0.006 | 0.013 |

*Live virus focus reduction neutralization assay was carried out in Vero-E6 cells. Dose-dependent neutralization of SARS-CoV-2 variants by mAbs and their combination was measured to obtain mAb potencies expressed as µg/mL).*

**Table S4.**
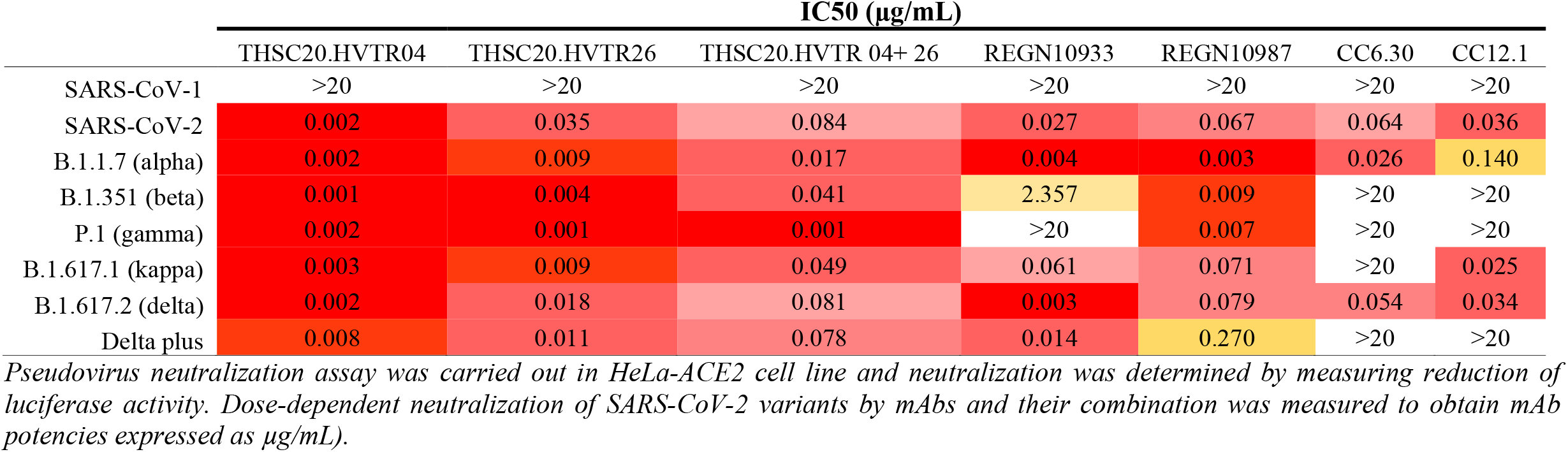
Neutralization breadth and potency of THSC20.HVTR04 and THSC20.HVTR26 and their combination against pseudoviruses expressing different SARS-CoV-2 VOC and VOI spike variants.

**Table S5.** Effect of key point mutations within RBM in SARS-CoV-2 spike on neutralization potency of THSC20.HVTR04 and THSC20.HVTR26 mAbs.

| *Mutations* | *Region* | **THSC20.HVTR04** |  | **THSC20.HVTR26** |  |
| --- | --- | --- | --- | --- | --- |
|  |  | *IC<sub>50</sub>* | *Fold reduction*<sup>§</sup> | *IC<sub>50</sub>* | *Fold reduction*<sup>§</sup> |
| WT (SARS-CoV-2) | - | 0.002 | - | 0.03 | - |
| R346K | RBD | 0.005 | 2.5 | 0.06 | 2 |
| K417N | RBD | 0.0015 | 0.75 | 0.05 | 1.66 |
| N439K | RBM | 0.3244 | 162.2 | 0.02 | 0.66 |
| N440K | RBM | 0.05 | 25 | 0.03 | 1 |
| K444N | RBM | 0.022 | 11 | 0.04 | 1.33 |
| F486L | RBM | 0.003 | 1.5 | 0.019 | 0.63 |
| F490S | RBM | 0.004 | 2 | 0.01 | 0.33 |
| N440K+D614G (B.1.36) | RBM/RBD | 0.182 | 91 | 0.02 | 0.66 |

*IC<sub>50</sub> values are given as µg/mL and were assessed in pseudovirus neutralization assay.*
<sup>§</sup> *Values represent fold reduction in IC<sub>50</sub> values when compared to the wild type (WT) SARS-CoV-2 spike in pseudovirus neutralization assay.*

### Supporting Figures

**Figure S1.**
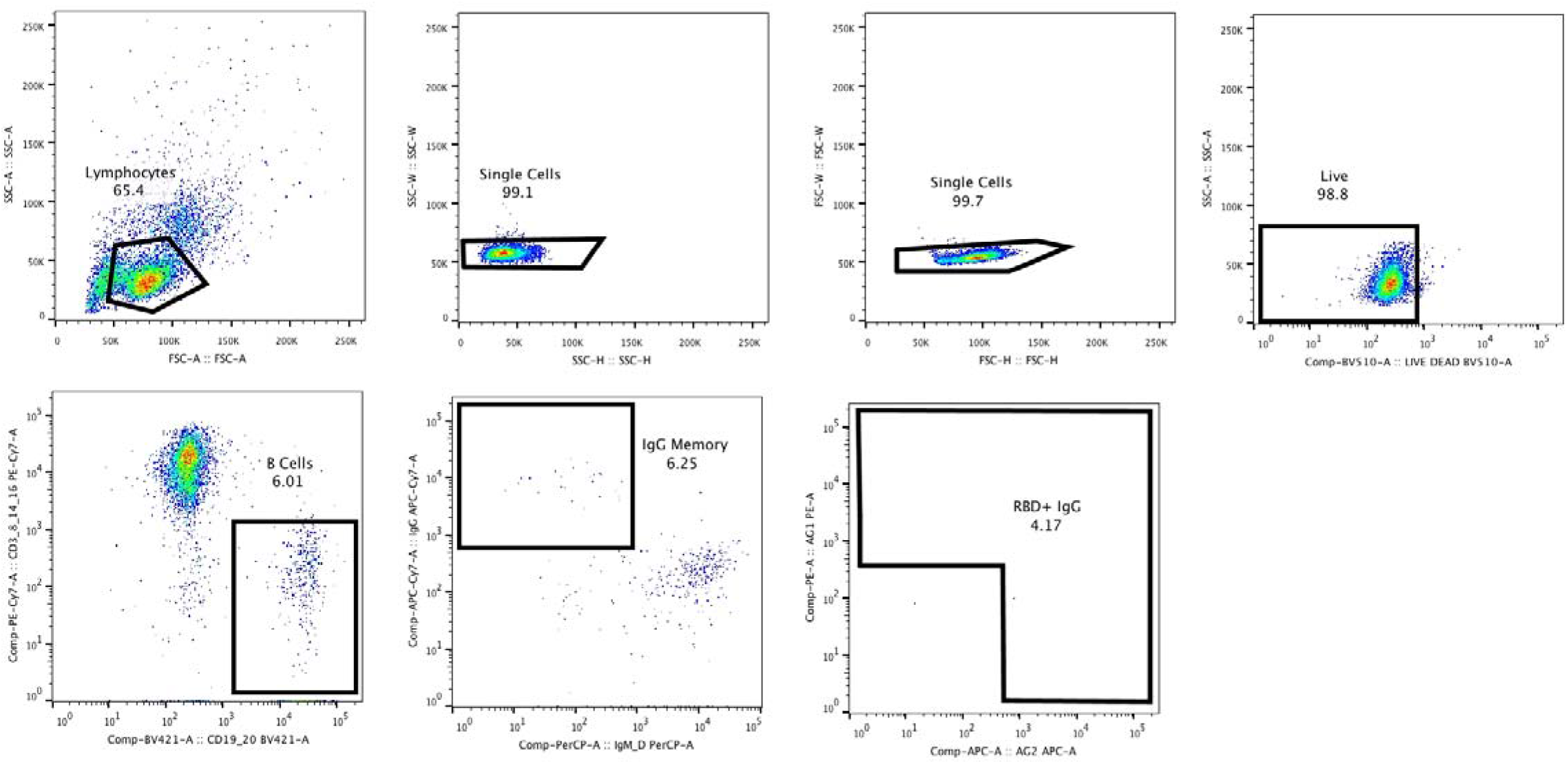
Representative Gating Strategy for the Antigen (RBD)-specific B cell sorting. Peripheral blood mononuclear cells (PBMCs) obtained from a convalescent donor C-03-0020 were stained with conjugated antibodies to cell surface markers, streptavidin labelled RBD probes and live dead stain and RBD-specific single B cells were sorted using a flow sorter. Singlet living CD19^+^C20^+^ IgG^+^ cells were gated and cells with positive SARS-CoV-2 RBD staining were selected for the single cell sorting into the 96 well plate prefilled with lysis buffer.

**Figure S2.**
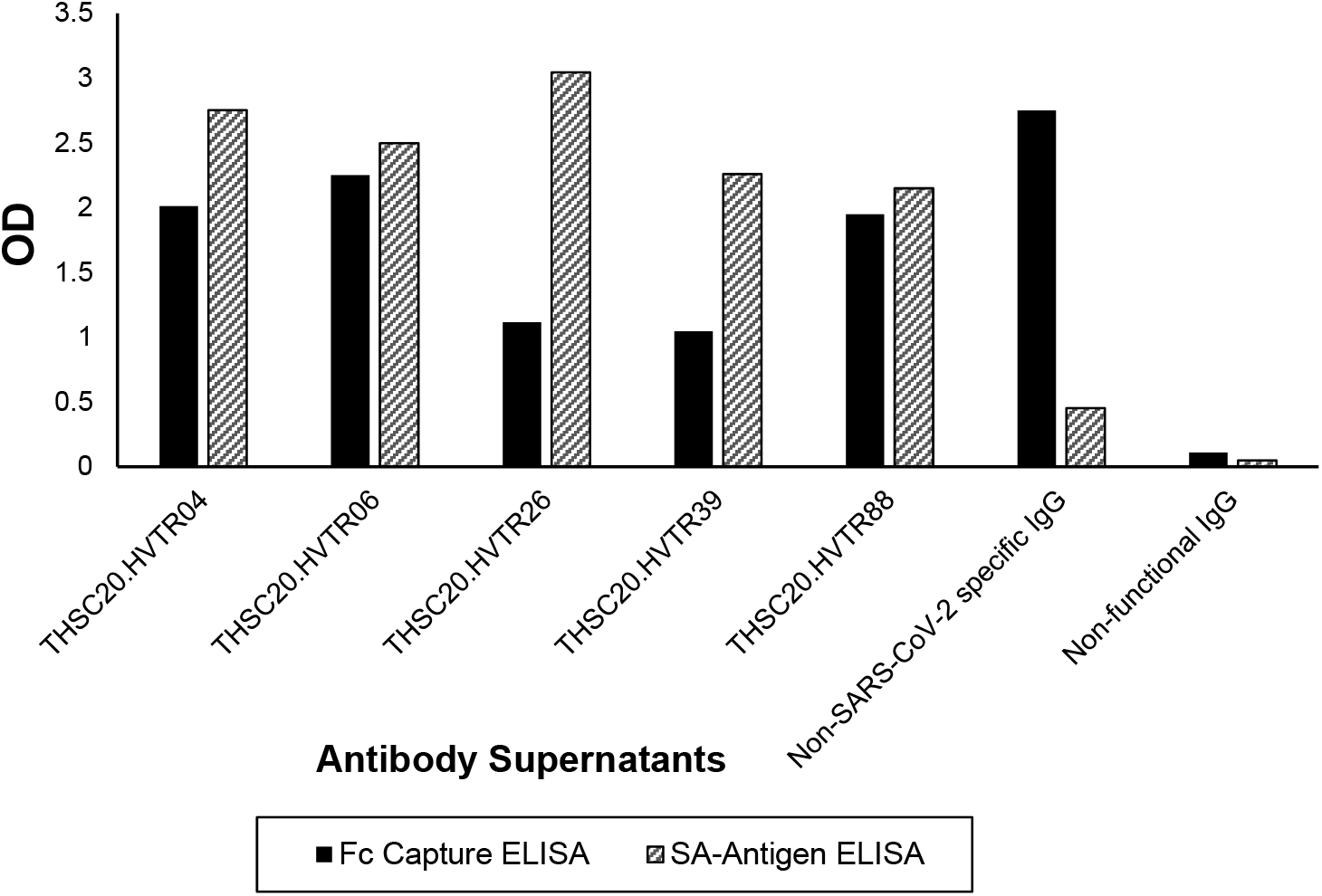
Expression and antigenicity of the isolated mAbs. Supernatants harvested from HEK 293T cells co-transfected with IgG expression vectors were examined for expression by Fc-capture ELISA (black filled bar) and their ability to bind to SARS CoV2 receptor binding domain (RBD) used for B cell sorting by streptavidin ELISA (striped line bar). Non-specific IgG refers to IgG that showed efficient expression but did not bind to RBD. Non-functional IgG refers to IgG sequence that neither expressed nor showed any RBD binding.

**Figure S3.**
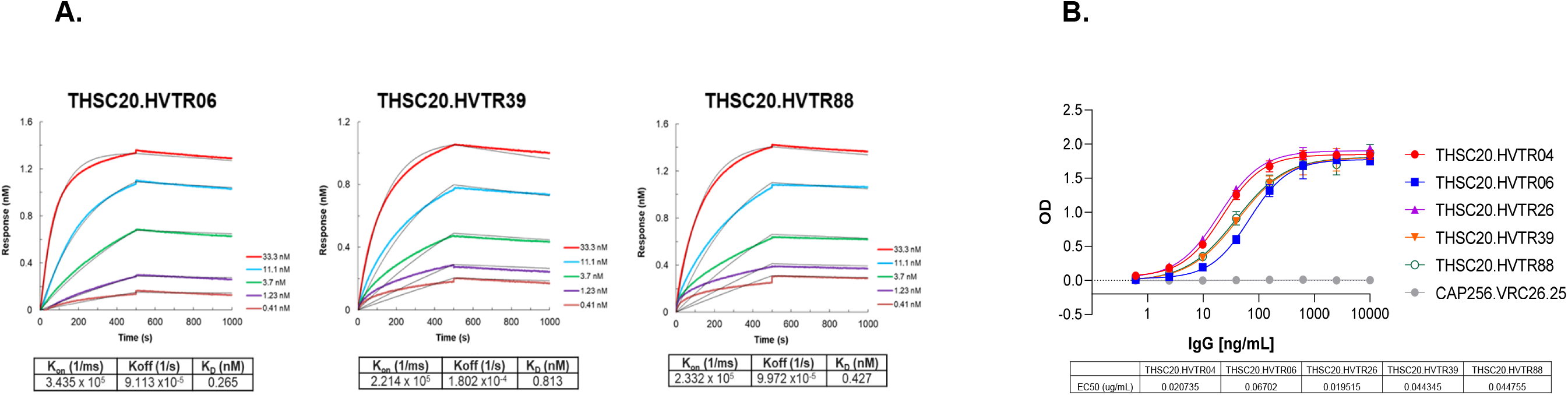
Binding affinity and avidity of five monoclonal antibodies with SARS-CoV-2 spike RBD by BLI and ELISA. A. Binding affinities of THSC20.HVTR06, THSC20.HVTR39 and THSC20.HVTR88 to the SARS-CoV-2 (Wuhan) receptor binding domain (RBD) protein by BLI-Octet. Biotinylated wild type SARS-CoV-2 RBD antigen was immobilized on Streptavidin (SA) biosensors and binding affinity of monoclonal antibodies to RBD was tested using three-fold serial dilutions of mAbs starting with 33.3 nM and lowest 0.41 nM (five different concentrations were tested). Association and dissociation was assessed for 500 seconds each. Data shown is reference-subtracted and aligned using Octet Data Analysis software v11.1 (Forte Bio). Curve fitting was performed using a 1:1 binding model and K_on_, K_off_ and K_D_ values were determined with a global fit. B. Binding avidity of mAbs determined by RDB-ELISA. Four-fold serial dilutions of mAbs starting with 10ug/mL were tested for binding to RBD by ELISA. Data shown mean with SEM from two replicates from single experiment. EC50 values were obtained by curve fit method using GraphPad Prism.

**Figure S4.**
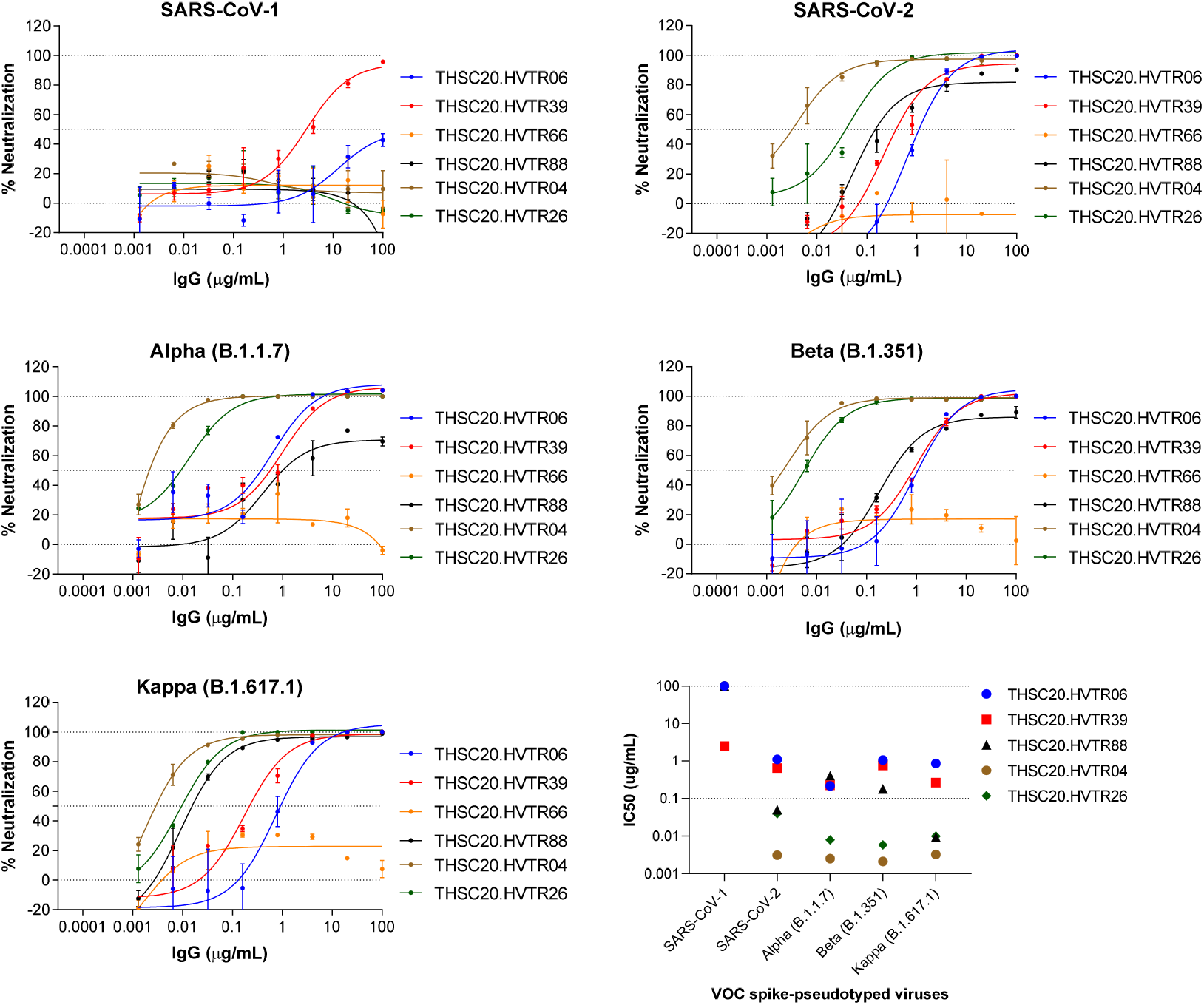
Comparison of neutralizing breadth and potency of all the mAbs isolated from the donor C-03-0020 in pseudovirus neutralization assay. Representative dose response curves from experiment with each concentration response tested in duplicate. THSC20HVTR04 and THSC20.HVTR26 mAbs were found to show maximum neutralization potency (lower panel, right) as determined by their IC50 values, obtained by non-linear regression four parameter curve fit method in GraphPad Prism. Shown values are mean with SEM.

**Figure S5.**
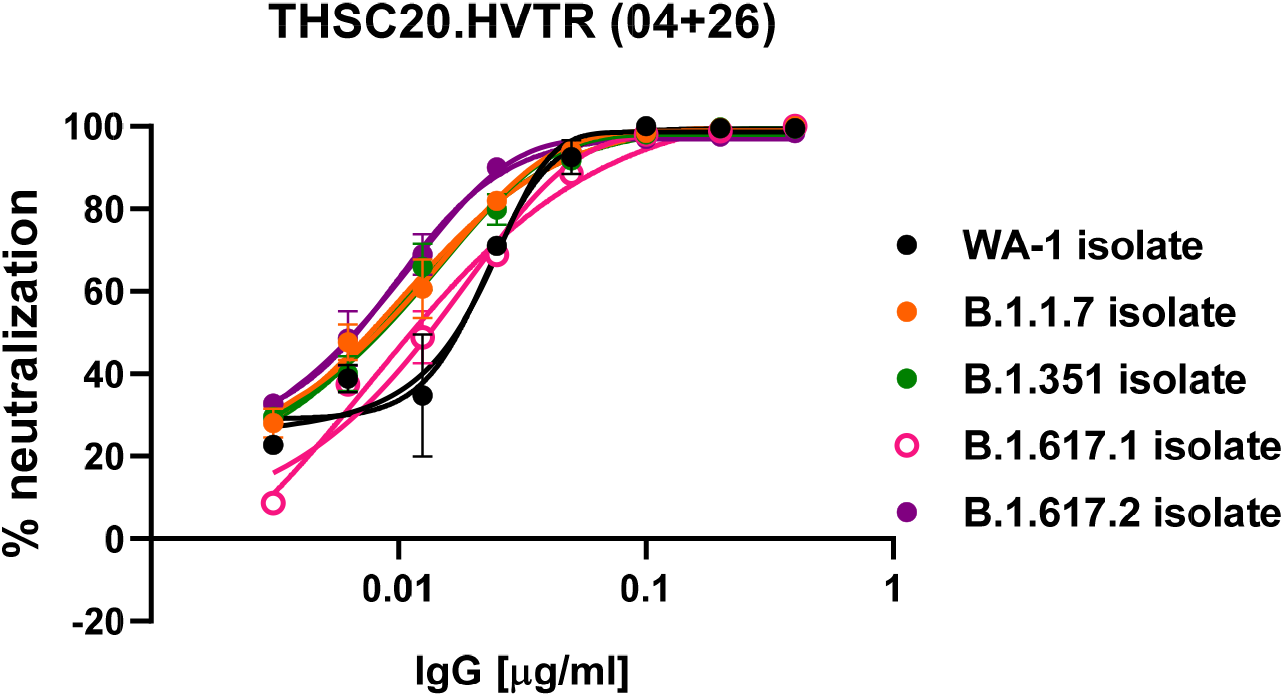
Live virus focus-reduction neutralization assay. The effect of combination of THSC20.HVTR04 and THSC20.HVTR26 was assessed by dose-dependent foci reduction neutralization (FRNT) live virus neutralization assay in Vero-E6 cells.

**Figure S6.**
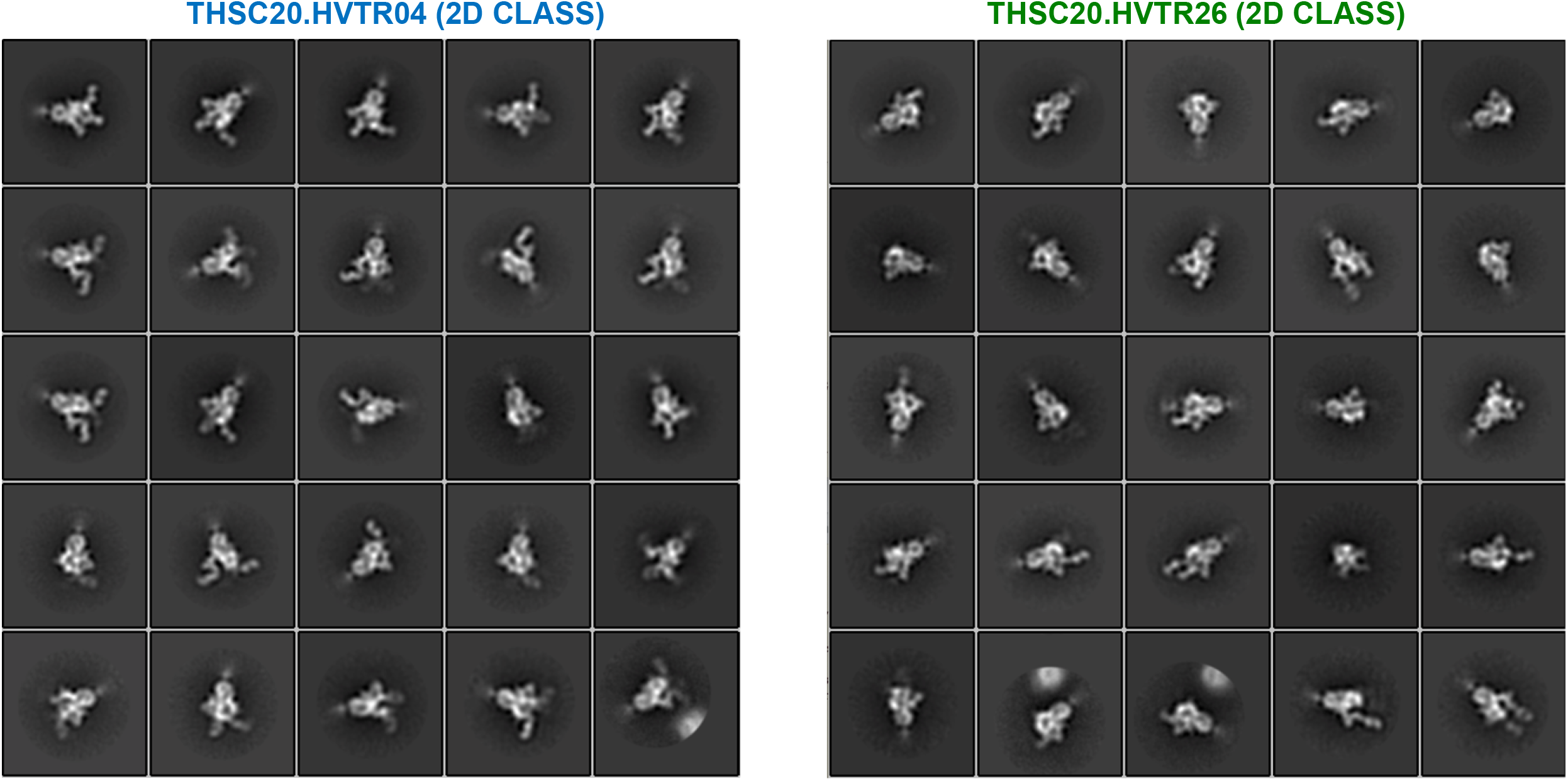
2D class averages of NS-EM of mAb-spike complexes. Low resolution images of the SEC purified complex of THSC20.HVTR04 and THSC20.HVTR26 Fabs with SARS-CoV-2 spike protein which were subsequently used for further refinement to generate high-resolution image as shown in Figure 3C.

**Figure S7.**
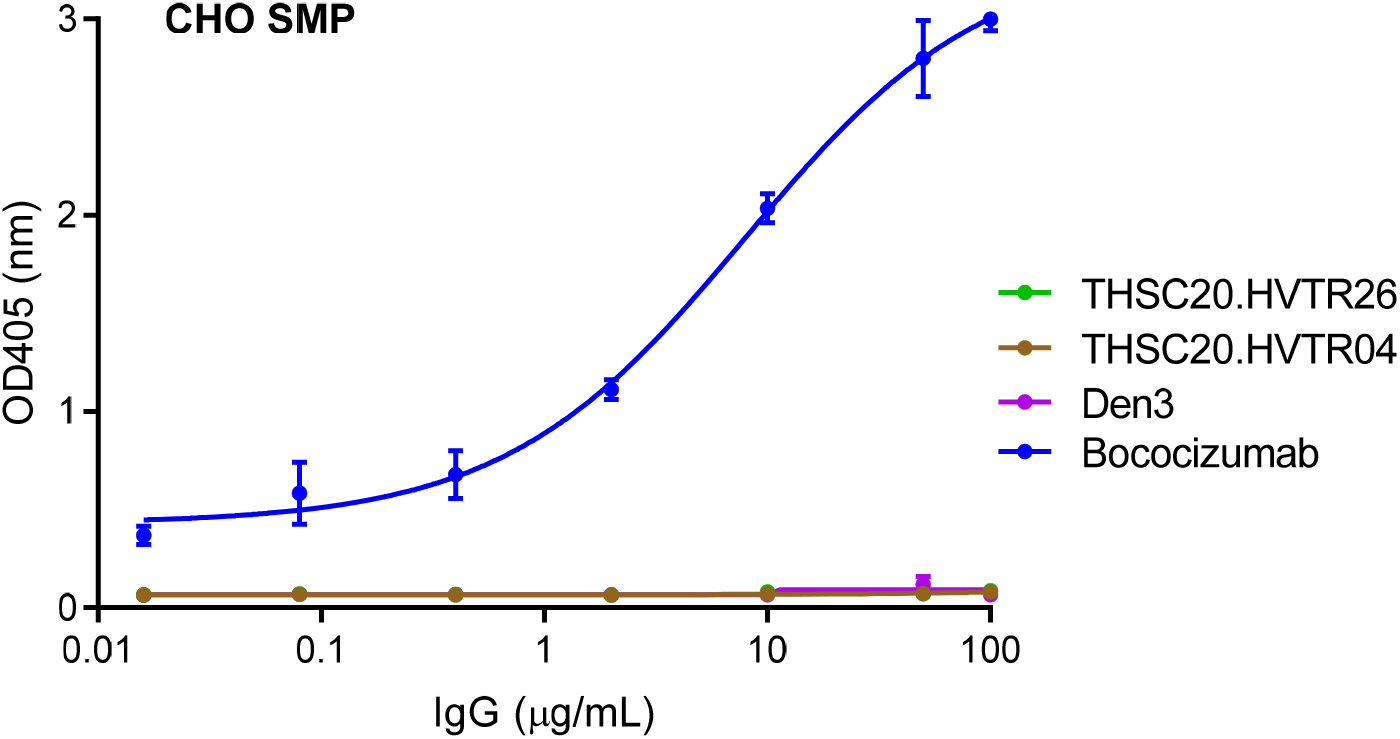
Poly-reactivity assessment of isolated mAbs. The polyreactivity of THSC20.HVTR04 and THSC20.HVTR26 mAbs using CHO soluble membrane protein (SMP) by ELISA. Three-fold serial dilutions of mAbs starting with 100ug/mL were tested.

**Figure S8.**
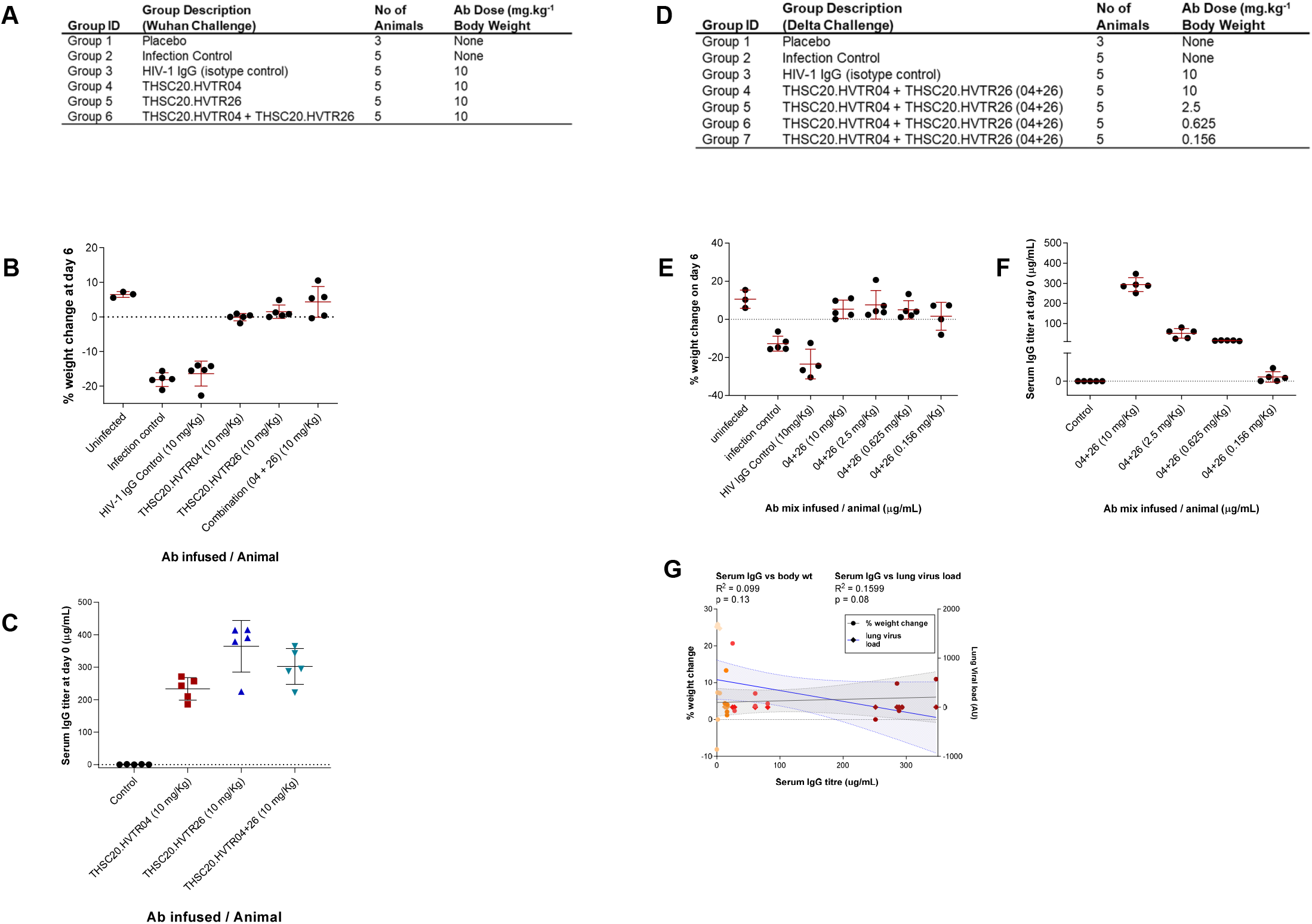
Dose-response effect of mAbs given singly and in combination on protection of mice against Wuhan and Delta infections. A. Animal grouping and antibody doses given (Wuhan challenge). B. Comparison of body weights between animals that received antibodies and those who did not prior to challenge with Wuhan isolate. C. Quantification of circulating serum IgG in mice at day 0 one day after infusion of mAbs at indicated doses and before virus (Wuhan) challenge. Values represent mean with SEM. D. Animal grouping and different antibody combination doses given (Delta challenge). E. Percent change in body weight of animals that received different doses of mAb combinations. Values represent mean with SEM. F. Quantification of circulating serum IgG concentration in mice at day 0 one day after infusion of mAbs at indicated doses and before virus (Delta) challenge. Values represent mean with SEM. G. Correlation between percent body weight change on day 6, circulating serum IgG concentration on day 0 and lung viral load on day 6 in mice those received different concentrations of mAb combinations.

